# Structural basis for emetine inhibition of ribosome translocation in *Toxoplasma gondii*

**DOI:** 10.64898/2026.07.23.740346

**Authors:** Wenzhao Dong, Fengrong Wang, Brenna A. Saladin, Suada Leskaj, Ritam Neupane, Vern Carruthers, Jailson Brito Querido

**Affiliations:** Department of Biological Chemistry, University of Michigan, Ann Arbor, MI 48109, USA; Life Sciences Institute, University of Michigan, Ann Arbor, MI 48109, USA; Center for RNA Biomedicine, University of Michigan, Ann Arbor, MI 48109, USA; Department of Microbiology and Immunology, University of Michigan Medical School, Ann Arbor, MI 48109, USA; Department of Biomedical Sciences, Iowa State University, Ames, IA 50010, USA

## Abstract

Apicomplexan parasites, including *Toxoplasma gondii* and *Plasmodium falciparum*, are major human pathogens that cause toxoplasmosis and malaria, respectively. The existing structures of *T. gondii* translational machinery are from empty ribosomes that lack several key components, including ribosomal protein RACK1 (Receptor for Activated C Kinase 1). Here, we used cryo-electron microscopy (cryoEM) to determine high-resolution structures of *T. gondii* ribosomal complexes, including a translating 80S ribosome bound to mRNA and tRNA. These structures reveal that RACK1 occupies the conserved binding site on the 40S subunit observed in other eukaryotic ribosomes. We also determined the architecture of the ribosomal P-stalk and identified the ribosomal proteins uL10 and uL11, which were not observed in previous *T. gondii* ribosome structures. In addition, we determined structures of the 80S ribosome bound to mRNA, tRNA, and the translation inhibitor emetine in two distinct conformational states. These snapshots reveal two mechanisms by which emetine inhibits the translocation step of mRNA translation: either by dislodging the mRNA from the E-site of the ribosome or by acting as a molecular glue within the E-site, thereby stalling translocation. Together, these findings provide new insights into the molecular basis of protein synthesis in apicomplexan parasites and establish a structural framework for the development of future antiparasitic therapeutics.

**GRAPHICAL ABSTRACT:** 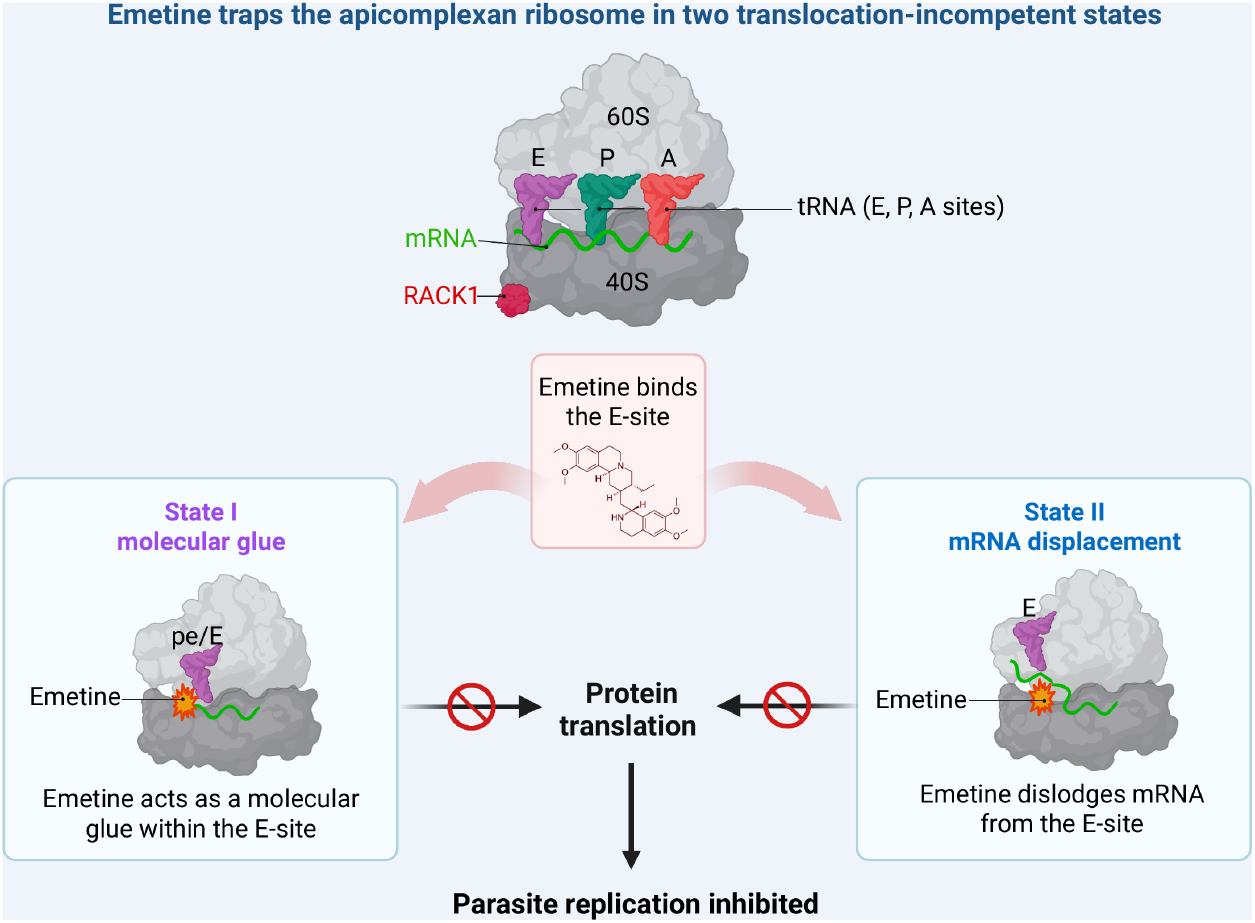

## INTRODUCTION

Protein synthesis is a key process for all domains of life and is carried out by the ribosome^1^. Thus, the ribosome is a major target for many antimicrobial and antiparasitic agents, as its central role in cell viability makes it an attractive point of intervention^2–5^. *Toxoplasma gondii*, an obligate intracellular parasite, infects approximately one-third of the world’s population^6^. Current treatments for acute *T. gondii* infection-primarily pyrimethamine and sulfadiazine are often limited by significant side effects and toxicity in some patients^7^. Moreover, there are currently no effective treatments available for eradicating the chronic bradyzoite stage^8^. Since translation is essential for *T. gondii* proliferation^9,10^, invasion of host cells^9,10,12^, and establishment of latent infection^11,13,14^, elucidating the structure of the ribosome in the parasite will be instrumental in development of next-generation therapeutics.

*T. gondii* possesses three distinct ribosomal systems: a eukaryotic-type cytosolic 80S ribosome, a highly divergent mitochondrial ribosome, and a prokaryote-like apicoplast ribosome^15–17^. Among these, the cytosolic 80S ribosome is responsible for the bulk of cellular protein synthesis, making it a key player in parasite biology and an attractive target for selective drug discovery.

Structural studies have been invaluable in advancing our understanding of ribosome function and its inhibition by small molecules^2,18,19^. High-resolution structures of the 80S ribosomes from diverse eukaryotic organisms, including yeast^20,21^, mammals^22,23^, and parasites^3,4,15,24,25^, have been determined by X-ray crystallography, cryo-electron microscopy (cryoEM), and cryo-electron tomography (in situ cryo-ET). For example, studies in *Plasmodium falciparum* have visualized the ribosomes in complex with tRNAs, translation factors, regulatory proteins, and small-molecule inhibitors^3,4,25^, revealing dynamic conformational states during translation and providing critical insights into mechanisms of action for ribosome-targeting drugs.

A prior cryoEM structure of the *T. gondii* 80S ribosome provided an important architectural foundation for understanding its unique features^15^. However, that structure captured the ribosome in an inactive state, lacking tRNAs, mRNA and translation factors, thereby limiting its functional interpretation. To address this gap, we used emetine, a classical-ribosome-targeting drug known to inhibit apicomplexan parasites by binding the small ribosomal subunit and inhibiting translocation^4^, to stall ribosomes in defined translational states. Using cryoEM, we captured and characterized distinct conformational states of *T. gondii* 80S ribosomes, including translating ribosomes, and an inactive ribosome bound to eukaryotic elongation factor 2 (eEF2).

Together, these structures expand our understanding of the functional dynamics of *T. gondii* 80S ribosome and provide a structural framework for future studies targeting translation in this parasite.

## RESULTS

### Structure of apicomplexan RACK1 bound to the ribosome

Human RACK1 and its yeast homolog Asc1 are highly conserved, multifunctional scaffold proteins that associate with the head of the 40S ribosomal subunit and play diverse roles in translation^26–30^. For example, RACK1 is required for efficient translation of some capped mRNAs, and broadens the translational capacity of the ribosome to accommodate a wider range of mRNAs^29^. The structures of RACK1 have been well characterized in both human ^31^ and yeast ^32^; however, the structure of *T. gondii* RACK1 remains unknown. To purify ribosomal complexes for cryoEM analysis, we perform sucrose density gradient ultracentrifugation to fractionate *T. gondii* cell extracts (Figure S1) derived from parasites cultured in fibroblasts. Prior to purification, we incubated *T. gondii* cell extracts with emetine to stall actively translating ribosomes. Mass spectrometry (MS) analysis of the purified 80S ribosome reveals the presence of ribosomal proteins, including RACK1 (TGME49_216880)(Supplementary table1). CryoEM analysis of the 80S fraction reveals distinct complexes, including an 80S ribosome containing mRNA and P-site tRNA (Figure 1A). When analyzing this *T. gondii* 80S structure, we identified a class of particles that contains additional density at the RACK1 binding site on the head of the 40S subunit. Focused classification on this region allowed us to improve the local resolution and unambiguously assign it as RACK1 (Figure 1B). The resulting density map, resolved to 3.0 Å, enabled us to build an atomic model that includes the small and large subunits of the 80S ribosome, the P-site tRNA, and mRNA (Figure 1A). In comparison with known *T. gondii* 80S structures ^15^, our structure revealed several previously unknown *T. gondii* translational machinery elements, including P-site tRNA, RACK1, as well as mRNA.

**Figure 1:**
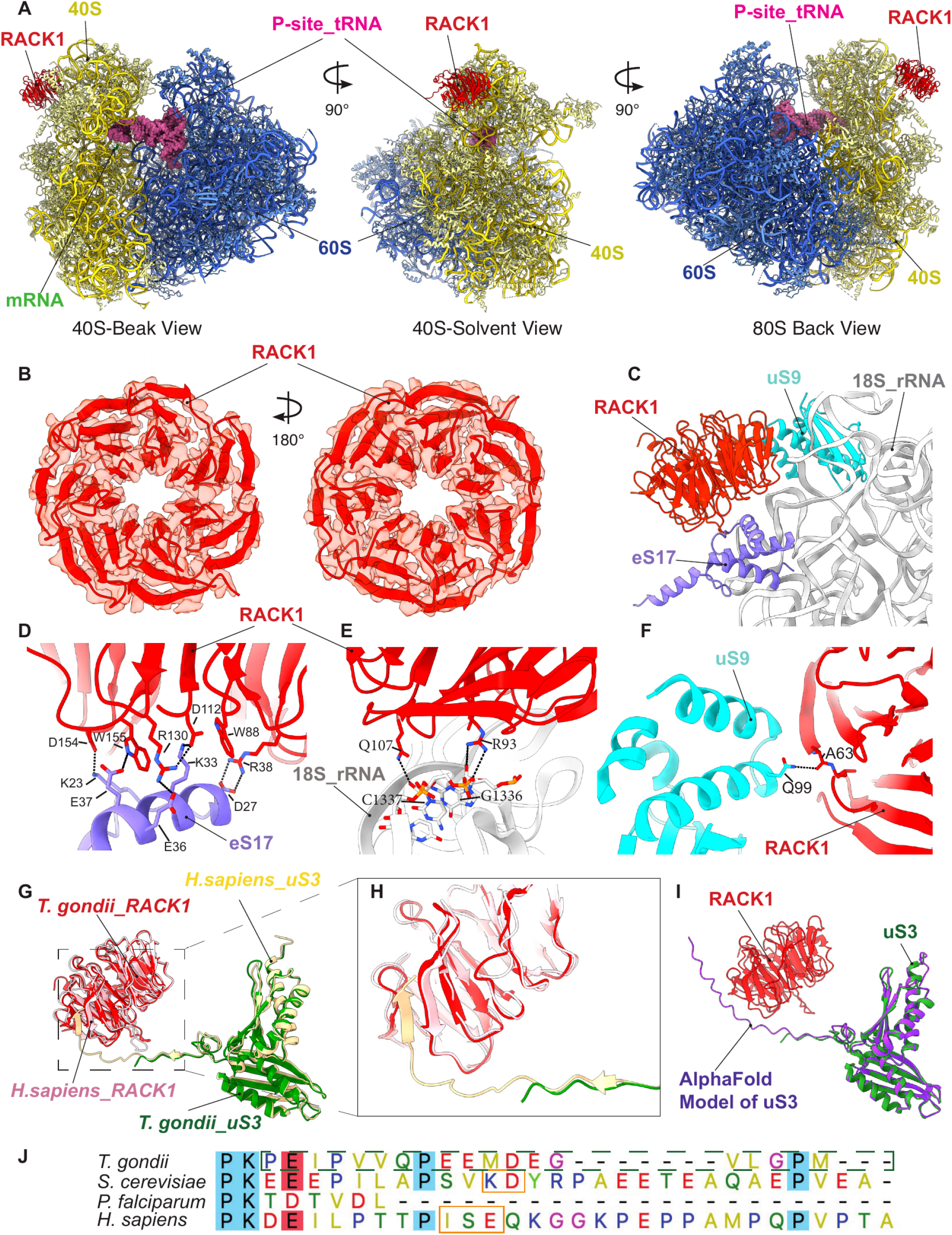
Overview of high-resolution structure of apicomplexan RACK1 bound to the 80S Ribosome. **(A** Molecular model of the 80S ribosome containing RACK1, P-site tRNA, and mRNA, shown in 40S beak view, 40S solvent view, and 80S back view. **(B** Molecular model of RACK1 fitted into the cryo-EM density map.**(C** Overview of RACK1 interactions with the surrounding ribosomal proteins eS17 and uS9, as well as 18S rRNA. **(D–F** Close-up views of the interactions between RACK1 and eS17 (D), RACK1 and 18S rRNA (E) and RACK1 and uS9 (F). **(G–H** Superposition of RACK1 and uS3 from *H. sapiens* and *T. gondii*, revealing that the interaction between RACK1 and uS3 observed in humans is absent in *T. gondii*. **(I** Superposition of uS3 from our model and the AlphaFold-predicted model. **(J)** Sequence alignment of RACK1 from *T. gondii* (A0A0F7UXI8), *S. cerevisiae* (P05750), *P. falciparum* (Q8IKH8), and *H. sapiens* (P23396). Alignments were generated using Muscle with defaults, Jalview. The region forming a β-sheet in yeast and human is highlighted by the orange box, while the region not observed in the *T. gondii* cryo-EM density map is indicated by the dotted green box.

Compared to human and *P. falciparum, T. gondii* RACK1 is conserved, with sequence identities of 61.2% and 67.3%, respectively (Figure S2A-B). In our model, RACK1 interacts with three components of the 40S small subunit: ribosomal RNA 18S, ribosomal protein uS9, and ribosomal protein eS17 (Figure 1C). These interactions are highly conserved in other eukaryotes such as humans and *P. falciparum*^31,33^. RACK1 primarily interacts with eS17 through eS17 residues Lys23, Asp27, Lys33, Glu36, and Glu37 (Figure 1D). At this interface, RACK1 residue Asp154 forms a hydrogen bond with eS17 Lys23, while RACK1 Arg38 forms a hydrogen bond with eS17 Asp27. Additional hydrogen bonds are formed between RACK1 residues Arg130, Asp112, and Trp155 and eS17 residues Lys33, Glu36, and Glu 37. Furthermore, RACK1 Trp88 engages in a π– cation interaction with eS17 Lys33. RACK1 further stabilizes its association with the ribosome through interactions with the 18S rRNA and uS9. Specifically, RACK1 residues Arg93 and Gln 107 contact the phosphate backbone of 18S rRNA nucleotides G1336, C1337 (Figure 1E), while RACK1 Ala63 interacts with uS9 residue Gln99 (Figure 1F).

In humans, the C-terminal extension of ribosomal protein uS3 interacts tightly with RACK1 through an antiparallel β-sheet, which plays a key role in the stabilization of RACK1 on the ribosome (Figure 1G-H)^31^. Notably, *T. gondii* uS3 does not interact with RACK1 (Figure 1G-I), indicating a significant divergence in ribosomal architecture between *T. gondii* and its mammalian host (Figure 1J). The structure of *T. gondii* uS3 is similar to that of *P. falciparum*^33^, both belong to apicomplexan parasites (Figure S2C). RACK1 is absent from most reported structures of apicomplexan ribosomes^4,15,34^, leading to the hypothesis that its loss reflects the presence of a truncated uS3 C-terminal tail that cannot reach RACK1. In *Plasmodium*, uS3 is indeed shorter (Figure 1J), consistent with this model. In contrast, *T. gondii* uS3 contains a sufficiently extended C-terminal tail to contact RACK1, as predicted by the AlphaFold model (Figure 1I). However, we do not observe any corresponding density, indicating that this region is conformationally flexible and does not bind RACK1. Together, these observations suggest that uS3 tail length alone is insufficient; productive association likely requires formation of specific antiparallel β-sheet interactions between the uS3 tail and RACK1.

### Structural basis for mRNA decoding in T. gondii

Translocation is a key step during translation elongation in which 80S ribosome moves along the mRNA, allowing tRNAs to move sequentially through the A-, P- and E-sites. We resolved three distinct ribosomal states by focused 3D classification of the tRNA-binding region. The first state contains a single P-site tRNA (Figure 2A and D). The second state contains a classical A-site tRNA(A/A) together with a hybrid P/E-site tRNA and corresponds to the RPE-translocation-hybrid 1 (PRE-H1) complex^35^ (Figure 2B and E). The third state also contains a classical A-site tRNA(A/A*) and a hybrid P/E-site tRNA but exhibits a subtle shift in tRNA position relative to PRE-H1 and corresponds to the PRE-translocation-hybrid 2 (PRE-H2) complex^35^ (Figure 2C and F). Together, these three structures capture successive early intermediates in the translocation pathway and provide direct structural evidence that these *T. gondii* ribosomes are actively engaged in translation.

**Figure 2:**
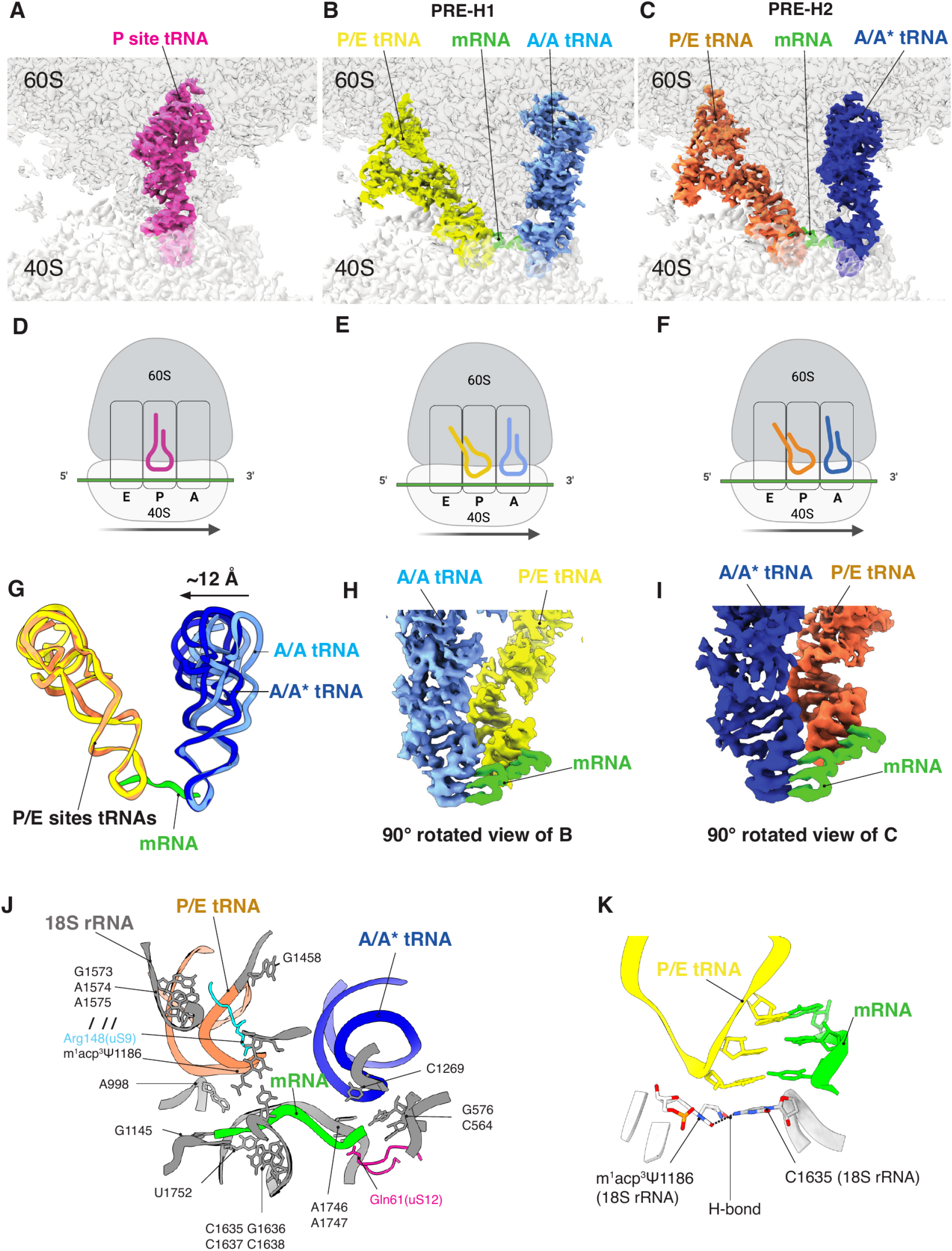
Structural basis of mRNA decoding and tRNA movement. **(A–C)** Overview of the cryo-EM electron density maps illustrating mRNA decoding and tRNA movement at three distinct stages: (A) a single tRNA positioned at the P site; (B) two tRNAs occupying the A/A and P/E sites; and (C) two tRNAs occupying the A/A* and P/E sites. **(D–F)** Corresponding cartoon models for panels A–C. **(G)** an ∼12 Å displacement of the peptidyl-tRNA elbow region for PRE-H1 and PRE-H2; **(H-I)** Close-up views of codon–anticodon interactions corresponding to panels B and C. **(J)** The interactions among mRNA, tRNAs and the 40S small ribosomal subunit; **(K)** m1acp3Ψ1186 interacts with the first anticodon nucleotide of the P/E-site tRNA and C1635 in the body domain of the 40S subunit.

For PRE-H1 and PRE-H2, both complexes contain small and large ribosomal subunits, showed densities for the peptidyl-tRNA in classical (A/A or A/A*) and the deacyl-tRNA in hybrid P/E states. These observations suggest that the complexes are in the pre-translocation (PRE) state, a transient intermediate in which tRNA–mRNA interactions are primed for translocation, although actual movement has not yet occurred^36^, which is consistent with the fact that these complexes lack eEF2. Our structure exhibits an intersubunit rotational movement of approximately 12° between the 40S small ribosomal subunit and the 60S large ribosomal subunit, which is consistent with the rotated-state conformations observed in other eukaryotes^35,37^. Furthermore, the two states revealed an ∼12 Å displacement of the peptidyl-tRNA elbow region, consistent with its transition from the canonical A/A state in PRE-H1 to the hybrid-like A/A* state in PRE-H2 (Figure 2G).

Closer inspection of the decoding center reveals a well-defined codon-anticodon interaction, with canonical base-pairing across the three nucleotide positions (Figure 2H-I). The mRNA–tRNA complexes are resolved at high resolution in the core region, allowing discrimination between purine and pyrimidine densities; however, unambiguous assignment to specific nucleotides remains challenging as the mRNA in this structure is endogenous and its sequence is unknown. The mRNA is stabilized through interactions with conserved components of the decoding center, including uS12 residues such as Gln61 and nucleotides within the 18S rRNA, including G1145, U1752, C1635, A1746 and G576. Similar to the mRNA, both tRNAs interact with multiple 18S rRNA nucleotides in a base-non-specific manner. For example, the peptidyl-tRNA contacts nucleotide C1269 of 18S rRNA, whereas the deacylated tRNA interacts with nucleotides A1574, A1575, G1573, and G1458 of the 18S rRNA, as well as Arg148 of ribosomal protein uS9. These interactions are highly conserved across eukaryotic species (Figure 2J)^35^.

The resolution also allowed us to observe one of the most prominent hyper-modifications, 1-methyl-3-α-amino-α-carboxyl-propyl pseudouridine 1186 (18S rRNA m^1^acp^3^Ψ1186; corresponding to Ψ1191 in yeast). This modification has been implicated in maintaining the architecture of the peptidyl-tRNA binding site^38^. In our structure, it is located within the head domain of the 40S small ribosomal subunit, where it interacts with the first anticodon nucleotide of the P/E-site tRNA. Simultaneously, a hydrogen bond is formed with the C1635 of 18S rRNA in the body domain of the 40S subunit, potentially stabilizing head-to-body interactions during translocation (Figure 2K).

### Structure of translationally inactive T. gondii eEF2-bound 80S complex

Eukaryotic elongation factor 2 (eEF2) is a GTPase that binds GTP and catalyzes its hydrolysis to drive ribosomal translocation ^21,39–43^. Additionally, eEF2 is well known for its activity as a hibernation factor^44–46^. While the structure of eEF2 has been elucidated in other species, including humans, the structure of *T. gondii* eEF2 remained elusive. Our cryoEM analysis reveals a class of 80S ribosomal particles that retain eEF2 but lack both mRNA and tRNA, consistent with a hibernating state (Figure 3A-C). eEF2 binding stabilizes the ribosome and enables us to determine the architecture of part of the P-stalk region, including densities corresponding to ribosomal proteins uL10 (also known as P0) and uL11 (Figure 3D), which were not observed in previous structures of the *T. gondii* ribosome^15^. eEF2 interacts with the ribosome through its interaction with ribosomal proteins uL10, uL11, and rRNA. These interactions are consistent with those observed in their mammalian host^47^.

**Figure 3:**
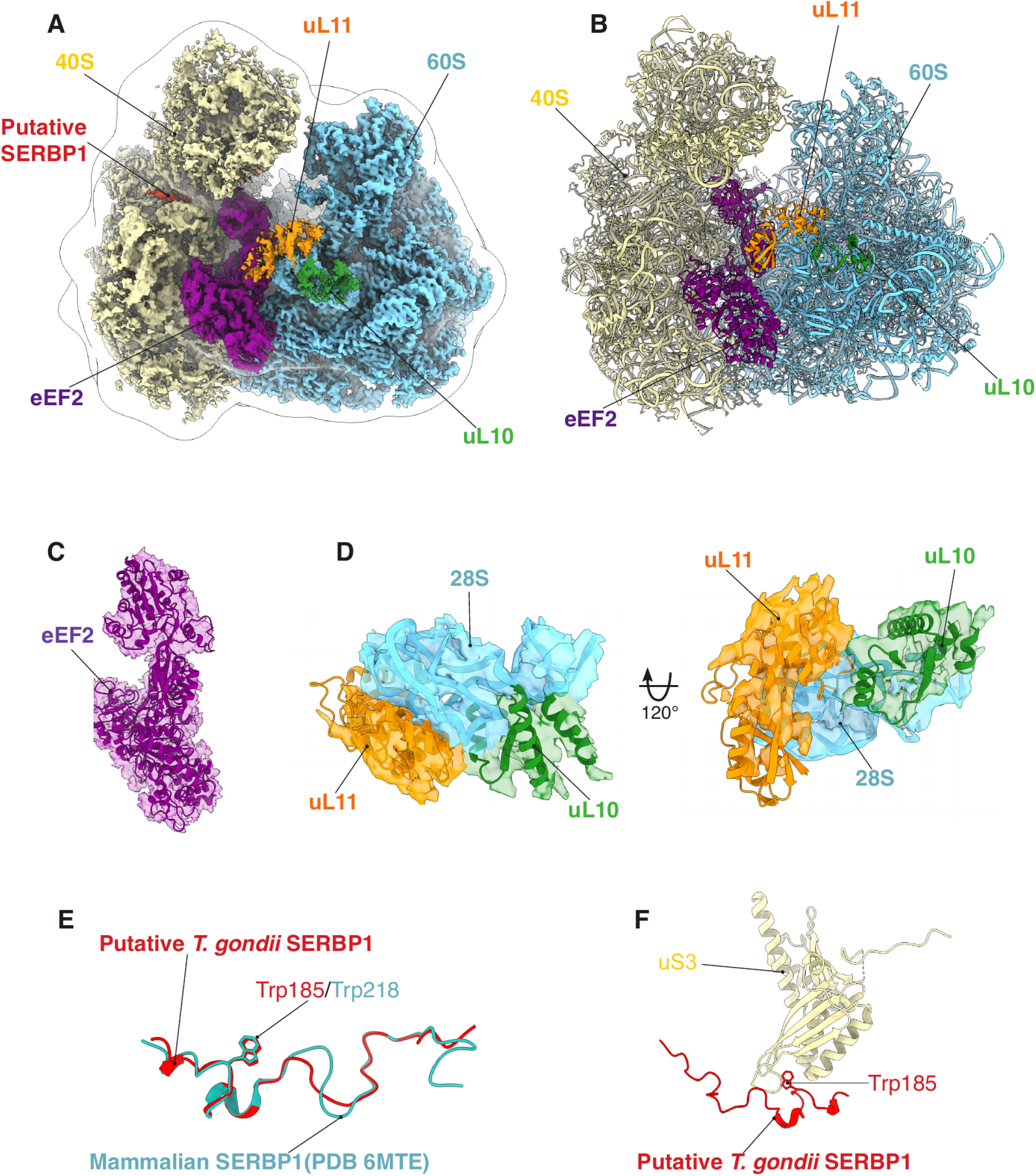
Structural overview of the hibernating *T. gondii* 80S ribosome. **(A–B)** Cryo-EM electron density map and corresponding molecular model of the hibernating *T. gondii* 80S ribosome, including eEF2, uL10, uL11, and a putative *T. gondii* SERBP1. Density corresponding to SERBP1 is observed in the mRNA channel but is not included in the molecular model as current resolution precludes unambiguous assignment. **(C)** Molecular model of eEF2 fitted into the cryo-EM density map. **(D)** Molecular model of uL10, uL11, and a portion of 28S rRNA fitted into the cryo-EM electron density map, highlighting interactions among these components. **(E)** Superposition of *T. gondii* SERBP1(SWISS-MODEL) and mammalian SERBP1(PDB 6MTE), atoms of Trp are shown in the models. **(F)** tryptophan residue binds to a hydrophobic pocket on uS3 within the mRNA channel.

Another key factor in ribosome hibernation is SERBP1 (serpine 1 mRNA-binding protein 1, also known as plasminogen activator inhibitor 1 RNA-binding protein), the mammalian homolog of yeast Stm1^45,48–51^. SERBP1 is conserved across most eukaryotes; however, no clear homolog has been identified in apicomplexan genomes, which are enriched in highly divergent, lineage-specific RNA-binding proteins^52^. This apparent absence has led to the hypothesis that apicomplexans use distinct factors to protect the mRNA channel of the 40S ribosomal subunit during ribosome hibernation. Notably, our structure reveals an unassigned density within the mRNA channel that resembles the density attributed to SERBP1 in other eukaryotic ribosome structures^45,48^. This density is also similar to that we recently described for the human tumor suppressor protein PDCD4 when bound to the mRNA channel^53^. However, we have not identified clear homolog of PDCD4 in *T. gondii* genome. Although the current resolution precludes unambiguous assignment, mass spectrometry and structural homology analyses identify TGME49_321680 (annotated as a putative RNA-binding protein) as the most likely *T. gondii* SERBP1 homolog, sharing ∼33% sequence identity with human SERBP1 (Figure S3). To generate an atomic model for structural comparison, we used SWISS-MODEL^54^ for homology modeling using mammalian SERBP1 as a reference^45^. A key binding partner of SERBP1 is the ribosomal protein uS3. SERBP1 Trp218 and PDCD4 Trp124 are highly conserved tryptophan residues that bind to a hydrophobic pocket on uS3 within the mRNA channel. We have previously shown that mutating this residue in PDCD4 abrogates the co-purification with the ribosome^53^. *T. gondii* TGME49_321680, present in our MS data, also contains this conserved residue (Figure 3E-F), consistent with its proposed homolog to human SERBP1.

To compare the *T. gondii* hibernating 80S ribosome with its human counterpart, we determined the structure of the human hibernating 80S ribosome, which has been reported previously^45,48^. Consistent with these earlier studies, our human structure contains a Z-site tRNA, eEF2, and the hibernation factors SERBP1 and EBP1^45,48^, but lacks mRNA (Figure 4A-B). Comparison of the human and *T. gondii* hibernating ribosomes revealed two major differences: the absence of Z-site tRNA and EBP1 in the *T. gondii* complex. One possible explanation is that, similar to RACK1, EBP1 associates more weakly with the *T. gondii* ribosome than with its human counterpart. In our human structure, EBP1 is stabilized by the ribosomal RNA expansion segment 27L (ES^27L^) (Figure 4C-E). To understand the potential structural differences, we used AlphaFold to predict the structure of *T. gondii* EBP1. The predicted structure shows substantial conservation between *T. gondii* and mammalian EBP1, particularly at putative ribosome-binding interface (Figure 4F). rRNA sequence alignment reveals substantial evolutionary divergence between *T. gondii* and its mammalian host^55,56^. ES^27L^ is the longest expansion segment in mammalian rRNA and is characterized by an extended GC-rich sequence. In contrast, ES^27L^ in *T. gondii* is considerably shorter and contains a lower GC content. These differences may weaken EBP1-ribosome interactions in *T. gondii* and contribute to the absence of EBP1density in our hibernating ribosome structure.

**Figure 4:**
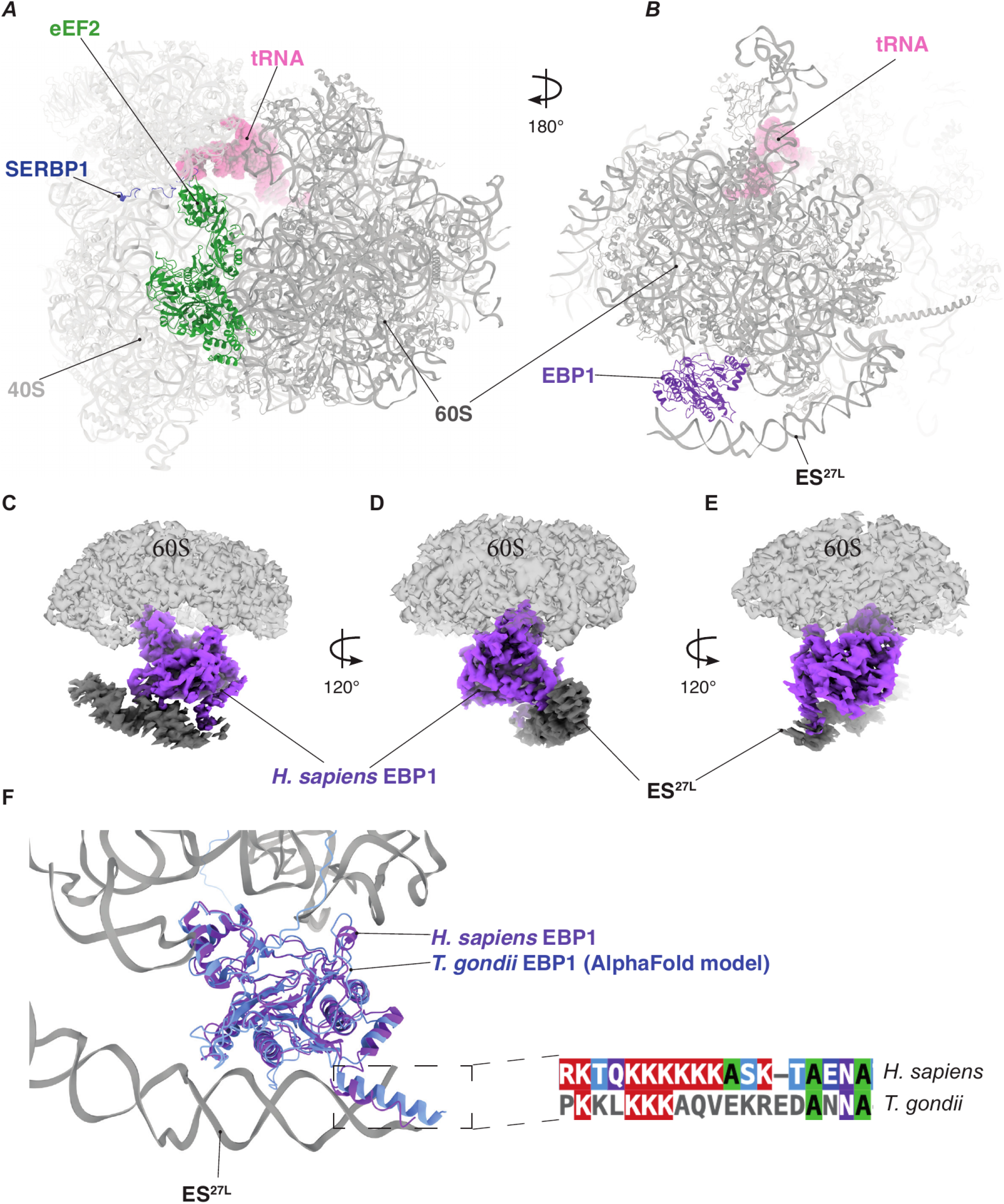
Structural overview of the hibernating human 80S ribosome. **(A-B)** Molecular model of the hibernating human 80S ribosome containing eEF2, tRNA, SERBP1, and EBP1. **(C-E)** Zoomed-in view of the density map showing EBP1 binding to ES27L **(F)** Structural superposition and C-terminal sequence alignment of *H. sapiens* and *T. gondii* EBP1(AlphaFold-predicted model), showing that *T. gondii* EBP1 tends to extend away from ES27L.

### Structure of T. gondii 80S bound to the antibiotic emetine

Emetine, a translation inhibitor, is a natural alkaloid that exhibits activity against a broad spectrum of protozoan parasites by inhibiting translocation during protein synthesis, including members of the phylum Apicomplexa, such as *P. falciparum* and *T. gondii* ^4,5,57,58^. An early cryoEM study mapped the emetine-binding site to the E site of the *P. falciparum* ribosome^4^. However, because that structure lacked both mRNA and tRNA, it did not provide the precise mechanistic insight into how emetine blocks translation. A more recent study sought to address this limitation by determining the structure of emetine bound to the *Giardia lamblia* ribosome^59^. In that case, ribosomal complexes were purified from cell extracts prior to drug incubation, a condition in which active translation is expected to be minimal or absent^60^. To overcome these limitations, we treated *T. gondii* cell extracts with emetine before isolating ribosomal complexes for cryoEM analysis (Figure S1), thereby capturing the drug-bound state in a more physiologically relevant context.

Our cryoEM classification revealed two distinct classes. In both classes, emetine occupies the E-site (Figure 5), consistent with previous observations in *P. falciparum* and *G. lamblia*^4,59^. Class 1 (C1) contains an 80S ribosome bound to emetine, mRNA, and a pe/E hybrid-state tRNA (Figure 5 A-C). Class 2 (C2) contains an 80S ribosome bound to emetine, mRNA, and an E-site tRNA in which the codon-anticodon interaction is dislodged from the mRNA channel in the 40S (Figure 5 D-F). The C2 class closely resembles the conformation reported for *G. lamblia*^59^, whereas the C1 class has not been described previously for emetine.

**Figure 5:**
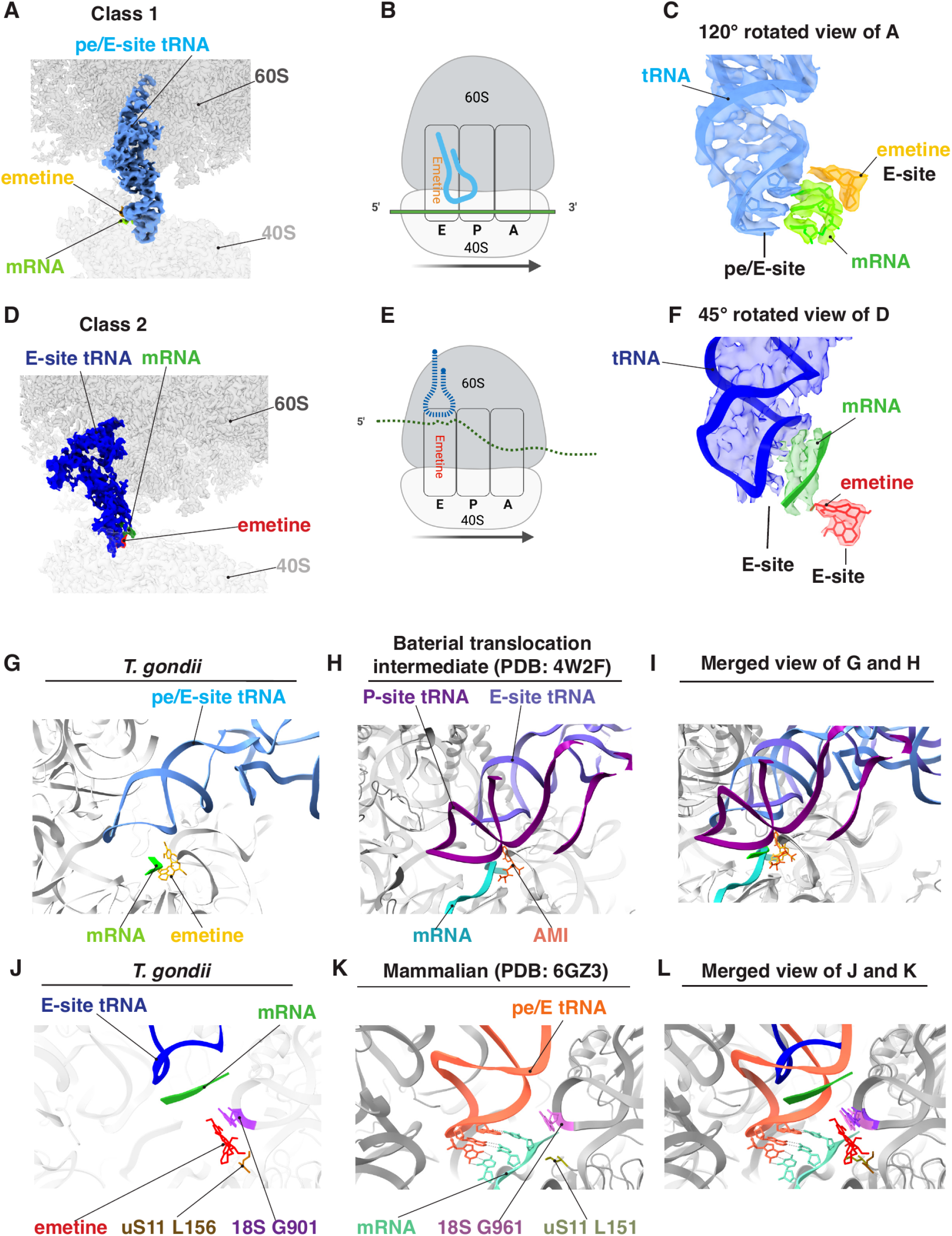
Structural basis of emetine binding to the *T. gondii* 80S ribosome. **(A)** Cryo-EM electron density map showing a tRNA positioned at the pe/E site and emetine bound at the E site. **(B)** Corresponding cartoon representation for panel A. **(C)** Close-up view of the molecular model fitted into the electron density map, highlighting the strong codon–anticodon interaction at the pe/E site and emetine binding at the E site. **(D)** Cryo-EM electron density map showing a tRNA positioned at the E site (displaced from the mRNA channel) and emetine bound at the E site. **(E)** Corresponding cartoon representation for panel A. **(F)** Close-up view of the molecular model fitted into the electron density map, highlighting the weak codon–anticodon interaction at the E site and emetine binding at the E site. **(G)**. Molecular model representation for panel A. **(H)**. Molecular model representation for bacterial translocation intermediate complexes stalled by the E site-binding antibiotic amicoumacin A(PDB 4W2F). **(I)**. Merged view of G and H. **(J)**. Close-up view of the *T. gondii* E site, showing upward displacement of the tRNA and mRNA relative to the canonical position, with emetine forming a base-stacking interaction with G901 of 18S rRNA and a hydrophobic interaction with L156 of uS17. **(K)** Close-up view of the mammalian E site (PDB 6GZ3), showing G961 contacts with tRNA and mRNA; the position of uS17 L151 is also highlighted. **(L)** Superposition of panels J and K, demonstrating that emetine occupies the E site and blocks the interaction between G901 and tRNA.

In the C1 class, the ribosome is captured in a translocation intermediate in which the anticodon stem-loop of a deacylated tRNA adopts a hybrid pe/E conformation between the P and E sites. This conformation resembles a bacterial translocation intermediate previously observed by X-ray crystallography in complexes stalled by the E site-binding antibiotic amicoumacin A^61^ (Figure 5 G-I). A similar translocation intermediate has also been reported in the presence of the eEF2 inhibitor, sordarin^35^. Together, these observations suggest that this conformation may represent a conserved translocation intermediate.

Consistent with prior studies that emetine binds within the ribosomal E-site, the drug is positioned adjacent to nucleotide G901 of *T. gondii* 18S rRNA, which occupies a position equivalent to G961 in mammalian ribosomes, a highly conserved residue implicated in stabilizing tRNA–mRNA interactions during translocation (Figure 5J–L)^62^. The benzo[a]quinolizine aromatic ring of emetine forms a π-stacking with G901, whereas the iso-quinoline ring interacts with the ribosomal protein uS11 Leu156, consistent with interactions reported in other organisms for both emetine and its analog cephaeline^4,59,63^. Structural comparison with a mammalian ribosome containing an accommodated E-site tRNA^62^ revealed that the anticodon stem-loop of the E-site tRNA is positioned immediately adjacent to G961 and uS17 Leu151 (Figure 5K). Overlay of the mammalian E-site tRNA with the *T. gondii* emetine-bound structure demonstrated that emetine occupies the same local environment surrounding G901 and uS11 Leu156 (Figure 5L). These observations suggest that emetine interferes with proper E-site tRNA accommodation and stabilization, thereby impeding the conformational rearrangements required for translocation.

### Emetine inhibits replication of acute-stage T. gondii tachyzoites

Because emetine binds the E-site of the *T. gondii* 40S ribosomal subunit, we hypothesized that it impairs parasite replication by inhibiting protein synthesis. To test this hypothesis, we assessed the effect o f e metine o n t he r eplication of acute-stage *T. gondii* tachyzoites. We allowed Type II ME49 Δ*ku80* tachyzoites to invade host cells for 3 h and then removed uninvaded parasites by washing. Next, we incubated the infected cells with either vehicle control or 300 nM emetine for 24 h. We quantified the number of parasites per vacuole as a measure of intracellular replication. Emetine treatment caused a significant shift in the distribution of vacuoles toward those containing fewer parasites compared with the control, indicating that emetine inhibits tachyzoite replication (Figure 6A,B).

**Figure 6:**
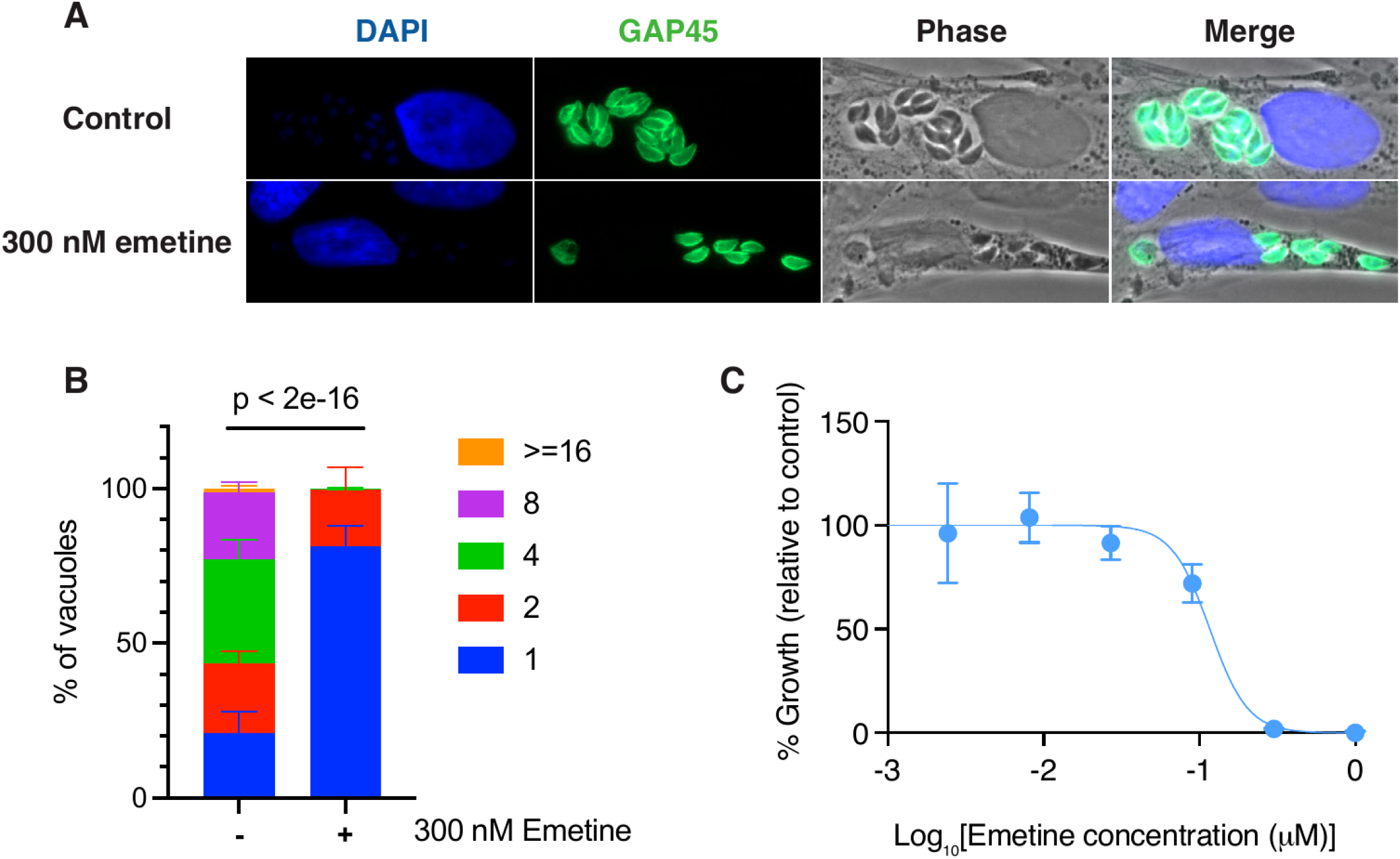
Biochemical basis of emetine-mediated inhibition. **(A-B)**. Emetine treatment inhibits replication of Type II *T. gondii* ME49 Δ*ku80* tachyzoites. **(C)**. Emetine inhibits replication of Type I *T. gondii* RH Δ*ku80* nLuc tachyzoites with an IC_50_ of 116.4 nM.

To further investigate the effects of emetine, we examined its impact on the growth of lab-adapted Type I *T. gondii* RH Δ*ku80* nLuc tachyzoites (Figure 6C). We allowed parasites to invade host cells for 4 h, removed uninvaded parasites by washing, and then incubated the infected cells with either vehicle control or increasing concentrations of emetine for 4 days. We quantified parasite growth using a NanoLuc luciferase assay. Emetine inhibited parasite growth in a dose-dependent manner, and nonlinear regression analysis estimated an IC_50_ of 116.4 nM (Figure 6C).

These findings are consistent with a recent study reporting that emetine inhibits the growth of both Type II Tg68 and ME49 tachyzoites (acute stage) with an EC_50_ of 30 nM^57^. However, the same study showed that emetine is less effective against in vitro bradyzoites (chronic stage), with EC_50_ values of 1.08 µM or 2.5 µM depending on the treatment used to induce differentiation i n v itro^57^. A dditionally, e metine demonstrated significant t oxicity t o h ost c ells ( EC_50_ = 2 30 n M t o 1.92 µM), which precluded further testing on ex vivo bradyzoites^57^. Together, these results demonstrate that emetine exerts robust inhibitory effects o n a cute-stage *T* . *g ondii* replication across multiple genetic backgrounds but suggest that derivatives of emetine with reduced host cell toxicity will be needed for further therapeutic development.

## DISCUSSION

Previous structural studies established the overall architecture of the *T. gondii* 80S ribosome but captured inactive particles lacking mRNA, tRNA, and associated regulatory factors, which limited mechanistic insight into translation in this parasite. In this study, we used emetine to trap ribosomes in defined functional states and determined multiple cryoEM structures that reveal translation, ribosome hibernation, and the mechanism of emetine-induced translation inhibition in *T. gondii*.

In this study, we present the structure of *T. gondii* RACK1 bound to the ribosome. RACK1 is a constitutive ribosomal protein in nearly all eukaryotes. However, early structures of *T. gondii* and *P. falciparum* ribosomes lacked this ribosomal protein^4,15,34^, raising questions about whether this protein associates with apicomplexan ribosomes. In *P. falciparum*, this question was recently addressed by two cryoEM studies^25,33^, whereas its association with *T. gondii* ribosomes has remained unresolved. Our structure reveals that RACK is a core structural component of the 40S small ribosomal subunit in *T. gondii*. Comparative analysis of RACK1 interactions with uS9, eS17, and 23S rRNA in human, yeast, *P. falciparum* and *T. gondii* reveals that these interactions are conserved, with the main difference being the interface between RACK1 and uS3.

RACK1 adopts a seven-bladed WD40 β-propeller architecture, in which each WD40 repeat is composed of four stranded antiparallel β-sheets. In humans and most eukaryotes, the C-terminal tail of uS3 (residues 223-226) forms an intermolecular antiparallel β-sheet interaction with WD40 repeat 4 of RACK1 (residues 183–189), which is critical for stabilizing RACK1 association with the ribosome. In *P. falciparum*, the C-terminal tail of uS3 is substantially shorter and lacks the region required for RACK1 binding, explaining the absence of RACK1 in most structural studies of the parasite ribosome. In contrast, although *T. gondii* uS3 is shorter than its human counterpart, it retains a sufficiently long C-terminal extension that could potentially interact with RACK1, as predicted by the AlphaFold model (Figure 1I). Nonetheless, our structure shows that the uS3 C-terminal tail does not bind RACK1, explaining the absence of RACK1 in previous structures of the *T. gondii* ribosome. Together, these observations indicate that formation of the intermolecular antiparallel β-sheet, rather than the length of the uS3 C-terminal tail itself, is the primary determinant for anchoring RACK1 to the 40S subunit. This model also provides a mechanistic explanation for why previous apicomplexan ribosome structures failed to detect RACK1.

Our structures also capture multiple active translation intermediates in *T. gondii*. We resolved ribosomes containing P-site tRNA as well as pre-translocation complexes containing A/A or A/A*- and P/E-site hybrid-state tRNAs (Figure 2). These structures reveal canonical codon-anticodon interactions within the decoding center and provide direct evidence that endogenous parasite mRNAs undergo active translation in our preparations. These structures reveal that despite extensive divergence in peripheral rRNA sequence, the molecular mechanisms that drive decoding and translocation remain strongly conserved in *T. gondii*.

The structure of the eEF2-bound *T. gondii* 80S ribosome provides important insight into ribosome hibernation and translational control in apicomplexan parasites (Figure 3). Although eEF2 classically functions as the essential GTPase that drives translocation during elongation, increasing evidence indicates that eEF2 also participates in translational repression and ribosome hibernation in eukaryotes. Our cryoEM structure indicates that *T. gondii*, similar to mammals and yeast, preserves conserved mechanisms that stabilize inactive ribosomes. The binding of eEF2 stabilized the ribosome and enabled us to resolve previously uncharacterized regions of the parasite ribosome, including portions of the P-stalk containing uL10 and uL11. The interaction network formed between eEF2, the P-stalk, and ribosomal RNA closely resembles those observed in other eukaryotes^47^, highlighting the remarkable conservation of this essential factor.

The structure also reveals a potential density for SERBP1 (Figure 3). In mammals, SERBP1 cooperates with eEF2 to stabilize inactive ribosomes and protect the mRNA channel during translational shutdown. Yeast Stm1 performs a similar function^48,49^, suggesting that hibernation factors represent a conserved strategy to preserve ribosomes during stress or nutrient limitation. However, no clear SERBP1 homolog had previously been identified in apicomplexan parasites, leading to the hypothesis that these organisms evolved entirely distinct mechanisms for ribosome hibernation. Structural comparison, mass spectrometry analysis, and sequence conservation strongly support TGME49_321680 as a likely functional homolog of SERBP1 in *T. gondii*. The conservation of this structural feature strongly suggests that apicomplexans preserve a common mechanism for protecting the mRNA channel during ribosome hibernation.

At the same time, our findings also highlight important divergence in the composition of hibernating ribosomes between *T. gondii* and its mammalian host. In contrast to the human hibernating 80S ribosome (Figure 4), we did not detect EBP1 or a Z-site tRNA in the parasite complex. EBP1 plays important regulatory roles in mammalian ribosomes and interacts extensively with the large-subunit expansion segment ES27L. Our structural and sequence analyses reveal that ES27L in *T. gondii* is substantially shorter and less GC-rich than its mammalian counterpart. This divergence likely weakens EBP1 association with the parasite ribosome and may explain its absence from our structure. These observations further suggest that expansion segments do not simply decorate the ribosome surface but instead actively shape the recruitment of regulatory factors during translational control.

A key advance of our study is the identification of two distinct emetine-stalled ribosome conformations, revealing that emetine inhibits translation through at least two mechanistically related states (Figure 5). In class C2, the E-site tRNA becomes displaced from the mRNA channel, disrupting codon–anticodon interactions. This conformation resembles that previously observed in *G. lamblia* and supports the model that emetine destabilizes proper E-site accommodation during translocation. However, class C1 has not been reported before. In this class, the ribosome was stalled in a translocation intermediate containing a deacylated pe/E hybrid-state tRNA positioned between the P- and E-sites. In addition to displacing the codon-anticodon interaction from the E site, emetine to trap the ribosome in an incomplete translocation state in which the tRNA cannot fully transition into the canonical E site.

The E site binders have been proposed to have two distinct mechanisms, one in which they displace the codon-anticodon from the E-site, such as that observed for pactamycin^61,63,64^, and an alternative mechanism in which they can both displace the codon-anticodon and act as a “glue” and prevent the codon-anticodon from fully entering into the E-site, resulting in a hybrid conformation, as described for amicoumacin A^61^. Our structure reveals that, similar to amicoumacin A, emetine can use two alternative mechanisms to stall translation. It is possible that the stalling mechanism is dictated by the nucleotide at positions -1 and -2 of the codon in the E-site. In this model, purines cannot be accommodated within the E site in the presence of emetine, resulting in codon-anticodon displacement and strong inhibition of translation, which may explain the previously suggested purine preference. On the other hand, pyrimidines can be partially accommodated into the E site even in the presence of emetine. In this case, it may result in a weak inhibition of translocation, with tRNA in a hybrid conformation that may have reduced affinity to eEF2.

This mechanism likely explains the potent inhibitory activity of emetine against acute-stage *T. gondii* tachyzoites observed in our replication assays (Figure 6). More broadly, these structures establish a framework for developing next-generation translation inhibitors that selectively stabilize transient ribosomal intermediates in apicomplexan parasites while minimizing toxicity toward the host ribosome.

## MATERIALS AND METHODS

### Emetine treatment of Type II T. gondii ME49 Δku80 tachyzoites

Host cells were human foreskin fibroblasts ( HFFs, Hs27; ATCC #CRL-1634), obtained from the American Type Culture Collection. The HFFs were cultured to confluence on coverslips in 6-well plates. Approximately 3 × 10^6^ *T. gondii* type II ME49 Δ *ku80*^1^ parasites were added to each well to infect the HFF monolayers. After 3 hours of invasion, unbound parasites were removed by washing the cultures three times with 1× PBS. Fresh D10 medium was then added, containing either control treatment (H_2_O) or 300 nM emetine (Sigma, 324693-250MG). The D10 medium was prepared using DMEM (Fisher Scientific; #10-013-CV), 10% heat-inactivated cosmic calf serum (Cytiva; #SH30087.03), 2 mM L-glutamine (Corning; #25-005-CI), and 50 U/mL penicillin-streptomycin (Gibco; #15070063). Cultures were maintained for 24 hours at 37°C with 5% CO_2_.

After incubation, the HFF monolayers were fixed with 4% formaldehyde for 10 minutes, permeabilized with 0.1% Triton X-100 in PBS for 10 minutes, and blocked with 10% FBS in PBS (containing 0.01% Triton X-100) for 30 minutes. Primary and secondary antibodies, as well as other staining reagents, were diluted in wash buffer (1% FBS, 1% normal goat serum, 0.01% Triton X-100 in PBS). Samples were incubated with primary antibodies overnight at 4°C, followed by a 1-hour incubation with secondary antibodies at room temperature. Rabbit anti-GAP45 primary antibodies^2^ (1:1000; Soldati Lab, University of Geneva) and goat anti-rabbit Alexa Fluor 488 secondary antibodies (1:1000; Invitrogen; #A11008) were used. After each step, samples were washed three times with wash buffer. Coverslips were mounted on slides with Mowiol (Calbiochem; #475904).

Images for quantification were captured on a Zeiss Axio Observer Z1 inverted microscope with a Plan Apochromat 63x/1.40 oil Ph3 M27 objective, using Zen Blue Edition (v2.6) software. All images within each experiment were acquired under identical imaging conditions. Parasites per vacuole were quantified by counting GAP45-positive parasites (GAP45 outlines each parasite) within each vacuole. Phase-contrast imaging was used to confirm vacuole identification. Each condition per biological replicate contained 189-305 vacuoles (from 30 images), with 3 independent replicates performed. Data represent mean ± s.d. Statistical significance was assessed using a generalized linear mixed model (GLMM) with a negative binomial distribution to account for overdispersion in parasite count data. The model was fitted using the glmer.nb function in the lme4 package in R. Drug treatment was included as a fixed effect, and biological replicate was included as a random intercept to account for variation between experiments. Statistical significance of fixed effects was evaluated using Wald z-tests. A P value < 0.05 was considered statistically significant.

### Emetine treatment of type I T. gondii RH Δku80 nLuc tachyzoites

HFFs were cultured to confluence in white, white-bottom 96-well plates (Falcon, #353296) and infected with 500 of type I *T. gondii* RH Δ*ku80* nLuc^3^ tachyzoites per well. Plates were incubated at 37°C with 5% CO_2_ for 4 hours to allow parasite invasion. After the incubation period, wells were washed once with PBS to remove unbound parasites, and fresh D10 media containing either emetine (Sigma, 324693-250MG) at various concentrations (1, 0.3, 0.09, 0.027, 0.0081, 0.00243, 0.000729, 0.000219, and 0.000066 µM) or H_2_O control was added. Plates were then returned to the incubator for an additional four days to allow parasite replication in the presence of the drug or control.

After four days of growth, parasite replication was assessed using a NanoLuc luciferase assay. Briefly, t he l uciferase substrate was prepared by diluting the stock solution to a final concentration of 12.5 μM in a 1:1 mixture of room-temperature PBS and 2× NanoLuc assay buffer ( 200 m M M ES, 2 mM CDTA, 1% Tergitol, 0.1% Antifoam 204, 300 mM KCl, 2 mM DTT, 70 mM Thiourea). After removing the media, 100 μL of the luciferase substrate was added to each well. Plates were incubated at room temperature for 10 minutes to allow cell lysis.

Luminescence was measured using a Synergy H1 plate reader (BioTek) with Gen5 software (Version 3.02) using the following settings: 1 s integration time; top, optics; gain of 135; 100 ms delay; and 4.5 mm read height. Data were normalized to H_2_O-treated control wells and expressed as percent growth. Dose–response curves were fitted by n onlinear r egression using the log(inhibitor) versus normalized response (variable slope) model in GraphPad Prism (v10.5.0). The half-maximal inhibitory concentration (IC_50_) was calculated from the best-fit curve. Data represent three independent biological replicates and are presented as mean ± s.d.

### Culturing parasites for harvesting 80S ribosome

Human foreskin fibroblasts (HFFs) were cultured to confluence in D150 dishes using D10 medium. Approximately 4×10^7^ *T. gondii* type II ME49 Δ*ku80*^1^ parasites were added per dish to infect the HFF monolayer. Infected cultures were maintained in D10 medium at 37°C with 5% CO_2_. At 1-2 days post infection, intracellular parasites were harvested by scraping the monolayers, followed by three passages through a 25-gauge needle, and filtering through a 3 μm Isopore™ membrane filter (Millipore; #TSTP02500). The filtered suspension was washed three times with cold PBS to remove cell debris. Parasites were pelted by centrifugation at 1,200 X g for 10 min.

The pellet was resuspended in lysis buffer containing 20 mM HEPES (pH 7.5), 1 mM DTT, 100 mM KCl, 3 mM MgCl_2_, EDTA-free protease inhibitor cocktail (Sigma; #P8340), RNase inhibitor (Promega; #N2611), and 3% glycerol. The suspension was flash-frozen i n l iquid n itrogen a nd s tored a t - 80°C until ready for 80S ribosome harvesting.

On the day of ribosome extraction, parasites were thawed, and Triton X-100 was added to the lysis buffer t o a final concentration of 0.1%. Lysates were incubated on ice for 10 min, and parasite lysis was monitored by microscopy. Complete lysis was confirmed by phase-contrast microscopy. Lysates were then centrifuged at 8,000 rpm for 5 min at 4°C to remove cell debris, and the supernatant was transferred to a new tube for subsequent ribosome purification. G MP-PNP a nd M gCl_2_ w ere a dded to final concentrations of 5 mM each, together with emetine to a final concentration of 50 μ M, and samples were incubated at 30°C for 10 min to stabilize actively translating ribosomes.

### Sucrose density gradient ultracentrifugation

Ribosomal complexes were separated using 10-30% (w/v) sucrose density gradients prepared in gradient buffer containing 20 mM HEPES-KOH (pH 7.5), 1 mM DTT, 100 mM KCl, and 3 mM MgCl_2_. Gradients were generated using a Gradient Master 108 (BioComp Instruments) in ultracentrifuge tubes (Seton Scientific) compatible with an SW41Ti rotor. Clarified lysates were loaded onto pre-cooled gradients and centrifuged at 40,000 rpm for 5 h at 4°C in a Beckman SW41 Ti rotor. Gradients were fractionated using a BioComp piston gradient fractionator with continuous monitoring at A260. Fractions corresponding to 80S ribosomes were identified from the absorbance profile and collected.

### Concentration and buffer exchange of 80S ribosomes

Collected 80S fractions were pooled and concentrated using Amicon Ultra centrifugal filters ( 10 k Da m olecular weight cutoff) pre-blocked with 1% BSA. Samples were concentrated by centrifugation at 4,251g at 4°C for 15 min. A 50 µl aliquot of the concentrated 80S sample (∼9.3 µg) was reserved for mass spectrometry analysis. The remaining 80S sample was washed repeatedly with cryo-EM buffer c ontaining 2 0 m M HEPES-KOH (pH 7.5), 100 mM potassium acetate, 4 mM magnesium acetate, 2% glycerol, 0.1 mM spermidine, 1 mM DTT, 50 μM emetine, and 0.2 U/μl RNAse by repeated centrifugation at 4,251g at 4°C to remove sucrose.

### Mass spectrometry

The 50 µl aliquot of the concentrated 80S sample (∼9.3 µg) was resuspended in 50 μl of 0.1M ammonium bicarbonate buffer (pH∼8). Cysteines were reduced by adding 50 μl of 10 mM DTT and incubating at 45° C for 30 min. Samples were cooled to room temperature and alkylation of cysteines was achieved by incubating with 65 mM 2-Chloroacetamide, under darkness, for 30 min at room temperature. An overnight (∼16 h) digestion with 1 ug sequencing grade, modified trypsin was carried out at 37° C with constant shaking in a Thermomixer. Digestion was stopped by acidification, and peptides were desalted using SepPak C18 cartridges using manufacturer’s protocol (Waters). Samples were completely dried using vacufuge. Resulting peptides were dissolved in 8 μl of 0.1% formic acid/2% acetonitrile solution and 2 μl of the peptide solution were resolved on a nano-capillary reverse phase column (Acclaim PepMap C18, 2 micron, 50 cm, ThermoScientific) using a 0.1% formic acid/2% acetonitrile (Buffer A) and 0.1% formic acid/95% acetonitrile (Buffer B) gradient at 300 nl/min over a period of 90 min (2-25% buffer B in 45 min, 25-40% in 5 min, 40-90% in 5 min followed by holding at 90% buffer B for 5 min and equilibration with Buffer A for 30 min). Eluent was directly introduced into Orbitrap Fusion tribrid mass spectrometer (Thermo Scientific, San Jose CA) using an EasySpray source. MS1 scans were acquired at 120K resolution (AGC target=2×10^5^; max IT=100 ms). Data-dependent High-energy C-trap dissociation MS/MS spectra were acquired using Top speed method (3 seconds) following each MS1 scan (NCE ∼32%; AGC target 5×10^4^; max IT 50 ms, 15K resolution).

Proteins were identified by searching the MS/MS data against a combined *Homo sapiens* and *Toxoplasma gondii* database using Proteome Discoverer (v3.0, Thermo Scientific). The database consisted of the reviewed *H. sapiens* protein database (20,347 entries; UniProt sp_canonical TaxID=9606, v2023-03-01, downloaded 05/10/2023) appended with the *T. gondii* ME49 protein database (8,316 entries; UniProt sp_tr_incl_isoforms TaxID=508771, v2023-09-13, downloaded 12/15/2023). Search parameters included MS1 mass tolerance of 10 ppm and fragment tolerance of 0.2 Da; two missed cleavages were allowed; carbamidimethylation of cysteine was considered fixed modification and oxidation of methionine, deamidation of aspergine and glutamine were considered as potential modifications. False discovery rate (FDR) was determined using Percolator and proteins/peptides with a FDR of ≤1% were retained for further analysis.

### Cryo-EM grid preparation, data acquisition and processing

3 µl of 80S ribosomes were applied onto glow-discharged UltraAUFoil Gold R1.2/1.3 grids (Quantifoil) pre-coated with a thin layer of graphene oxide (Sigma-Aldrich) made in-house. Cryo-EM grids were prepared using a Vitrobot (ThermoScientific) at 4°C and 100% humidity with a blot time of 8s and blot force of -15 and then plunged into liquid ethane at approximately 93 K.

The dataset was collected on a Titan Krios G4i (ThermoFisher) equipped with a Gatan K3 direct detector and a BioQuantum imaging filter at a magnification of 105,000X, resulting in a pixel size of 0.832 Å/pixel. For each exposure, 60 frames were collected with a dose of 1 e/Å^2 per frame. The target defocus range was set between -1.2 µm and -3.0 µm.

### Image processing

Micrographs were processed using Relion-5.0. Motion correction was carried out using Relion’s built-in implementation. Contrast Transfer Function (CTF) estimation was performed using CTFFIND4.1. The selected 2D classes were utilized for reference-based 3D classification. A cryo-EM map of a *T. gondii* 80S ribosome was used as a reference after low-pass filtering to 60 Å. Classes of particles exhibiting eEF2 or tRNA density were selected for Bayesian polishing in RELION to correct beam-induced motion. Following polishing and CTF refinement, mask classification was conducted focusing on the eEF2, RACK1 or the tRNA.

### Model building and refinement

We initiated model building and refinement by rigid-body fitting the prior structure of the Toxo ribosome and alpha fold predicted RACK1 or eEF2 into the cryo-EM map using Coot. Subsequently, we refined the model using Phenix for real-space refinement, followed by manual fitting to address any outliers.

### Cryo-EM figure generation

Figures were made using ChimeraX.

## ACKNOWLEDGEMENTS

We acknowledge the University of Michigan’s Proteomics Resource Facility (RRID: SCR_026723), for providing assistance with proteomic data acquisition. We thank the contributors to ToxoDB for invaluable resources to our work. We thank Biorender for providing a user-friendly tool for creating schematic illustrations. This work was supported by a grant from the US National Institute of Health, R01AI120607 (V.B.C.). Rogel Cancer Center (NIH Award Number P30 CA046592) (to J.B.Q.) and LSI computer cluster (S10OD030275) (to J.B.Q.). U-M cryo-EM Facility is supported by the U-M Life Sciences Institute, U-M Biosciences Initiative, and Beckman. American Heart Association (25CDA1450206 to F.W.) and the Knights Templar Eye Foundation (F.W.).

## AUTHOR CONTRIBUTIONS

J.B.Q., W.D., and F.W. conceived the study. W.D. and F.W. performed most of the experiments and data analyses. B.S., S.L. and R.N. assisted with data analysis. B.S. prepared and analyzed human 80S samples. J.B.Q. and V.B.C. supervised the study. W.D. and F.W. generated the illustrations. W.D., F.W., V.B.C., and J.B.Q. wrote the manuscript with input from all authors.

## DATA AVAILABILITY

The cryo-EM density maps and corresponding atomic coordinates for the structures reported in this study have been deposited in the Electron Microscopy Data Bank (EMDB) and the Protein Data Bank (PDB) under accession codes EMDB-YYY/PDB-XXXX (80S_RACK1_P-tRNA), EMDB-YYY/PDB-XXXX (80S_eEF2), EMDB-YYY/PDB-XXXX (PRE-H1), EMDB-YYY/PDB-XXXX (PRE-H2), EMDB-YYY/PDB-XXXX (80S-emetine Class 1), EMDB-YYY/PDB-XXXX (80S-emetine Class 2), and EMDB-YYY/PDB-XXXX (human hibernating 80S). The accession numbers will be updated upon publication. All other data supporting the findings of this study are included in the article, the Supplementary Information, or are available from the corresponding author upon request.

## Supplemental figures and tables

**Supplemental Figure 1:**
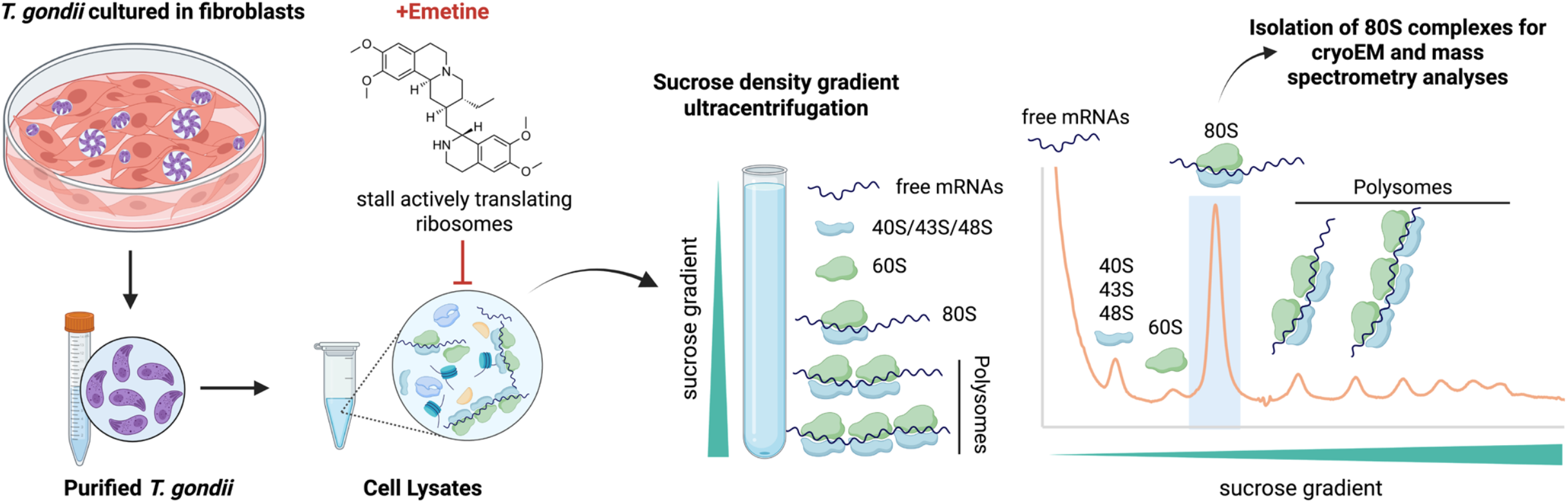
Purification workflow for actively translating *T. gondii* 80S ribosome complexes for cryoEM and mass spectrometry analyses. *T. gondii* parasites were cultured in human fibroblasts and purified prior to cell lysis. Emetine was added to stall actively translating ribosomes on mRNAs and preserve translation complexes. Cell lysates were subjected to sucrose density gradient ultracentrifugation to separate ribosomal species, including free mRNAs, 40S/43S/48S particles, 60S subunits, 80S monosomes, and polysomes. Fractions corresponding to 80S ribosome complexes were collected from the sucrose gradient and used for cryoEM and mass spectrometry analyses.

**Supplemental Figure 2:**
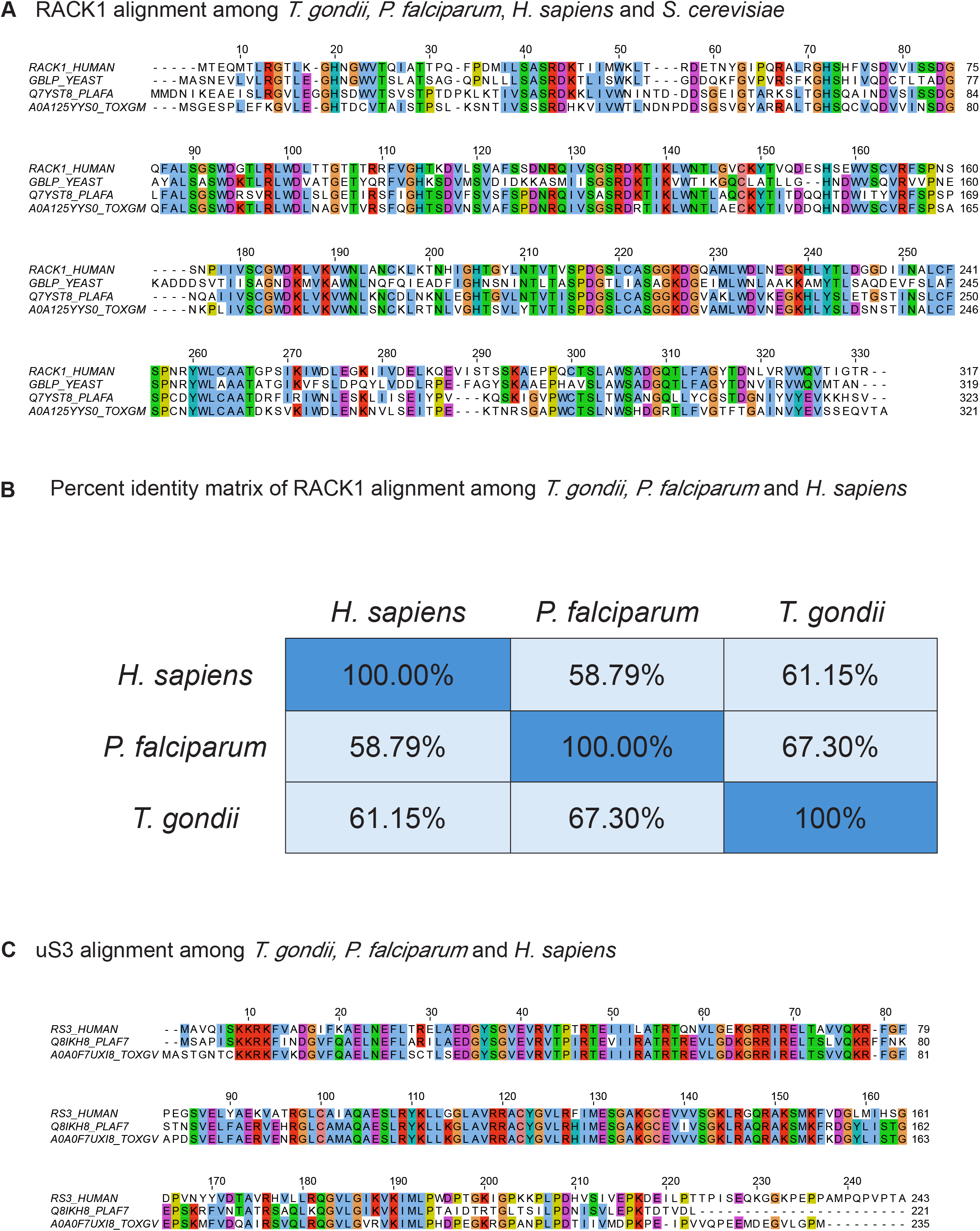
Sequence conservation of RACK1 and uS3 among *T. gondii, P. falciparum*, human, and yeast. **(A)**. Multiple sequence alignment of RACK1 proteins from *T. gondii, P. falciparum*, human and yeast. Conserved residues are highlighted with darker shading indicating higher sequence conservation. Alignments were generated using Muscle with defaults, Jalview. **(B)**. Percent identity matrix showing pairwise amino acid sequence identities *T. gondii, P. falciparum* and human. **(C)**. Multiple sequence alignment of uS3 proteins from *T. gondii, P. falciparum* and human. Conserved residues are highlighted with darker shading indicating higher sequence conservation. Alignments were generated using Muscle with defaults, Jalview.

**Supplemental Figure 3:**
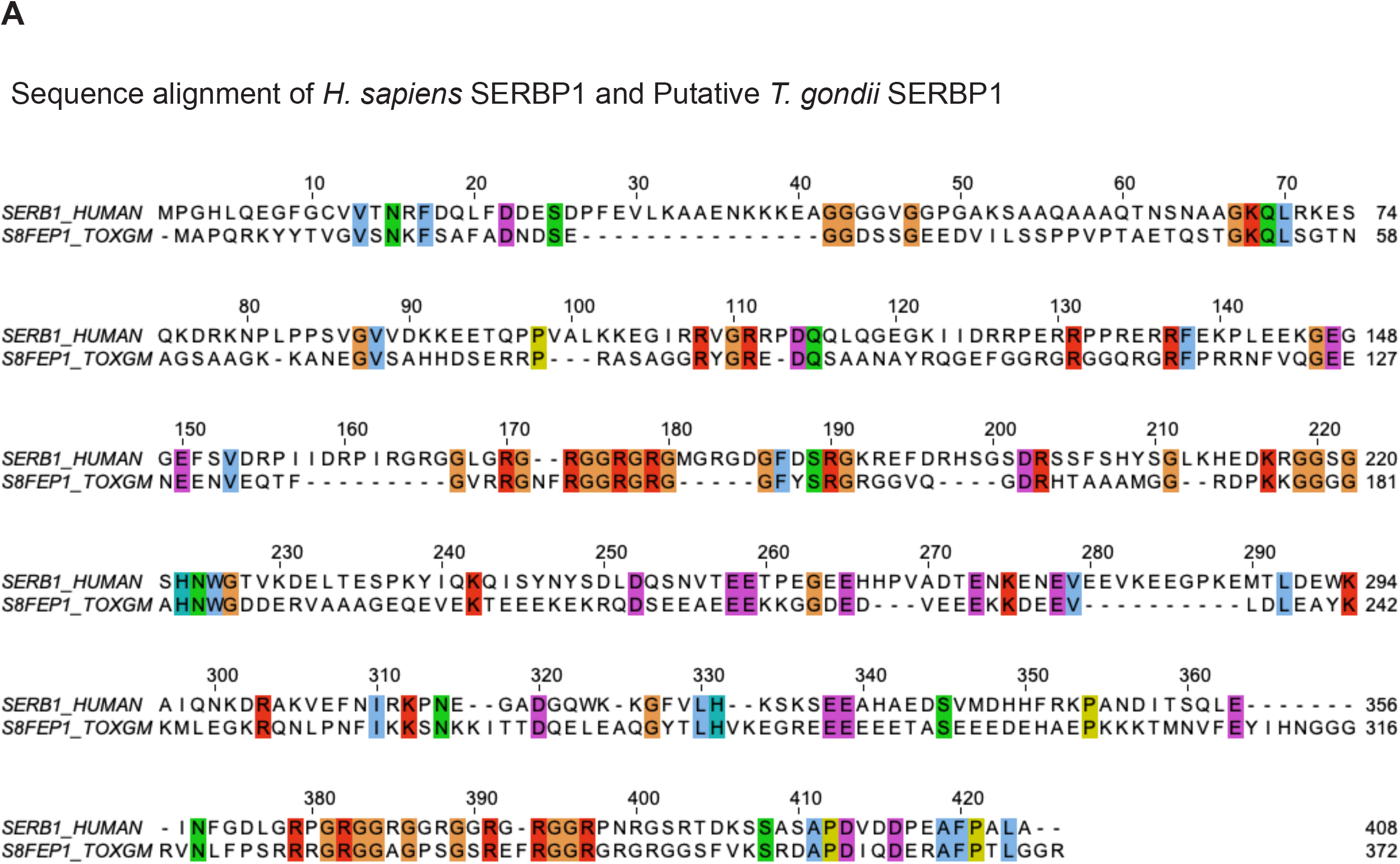
Sequence alignment of *H. sapiens* SERBP1 and Putative *T. gondii* SERBP1. Alignments were generated using Muscle with defaults, Jalview.

**Supplemental Figure 4:**
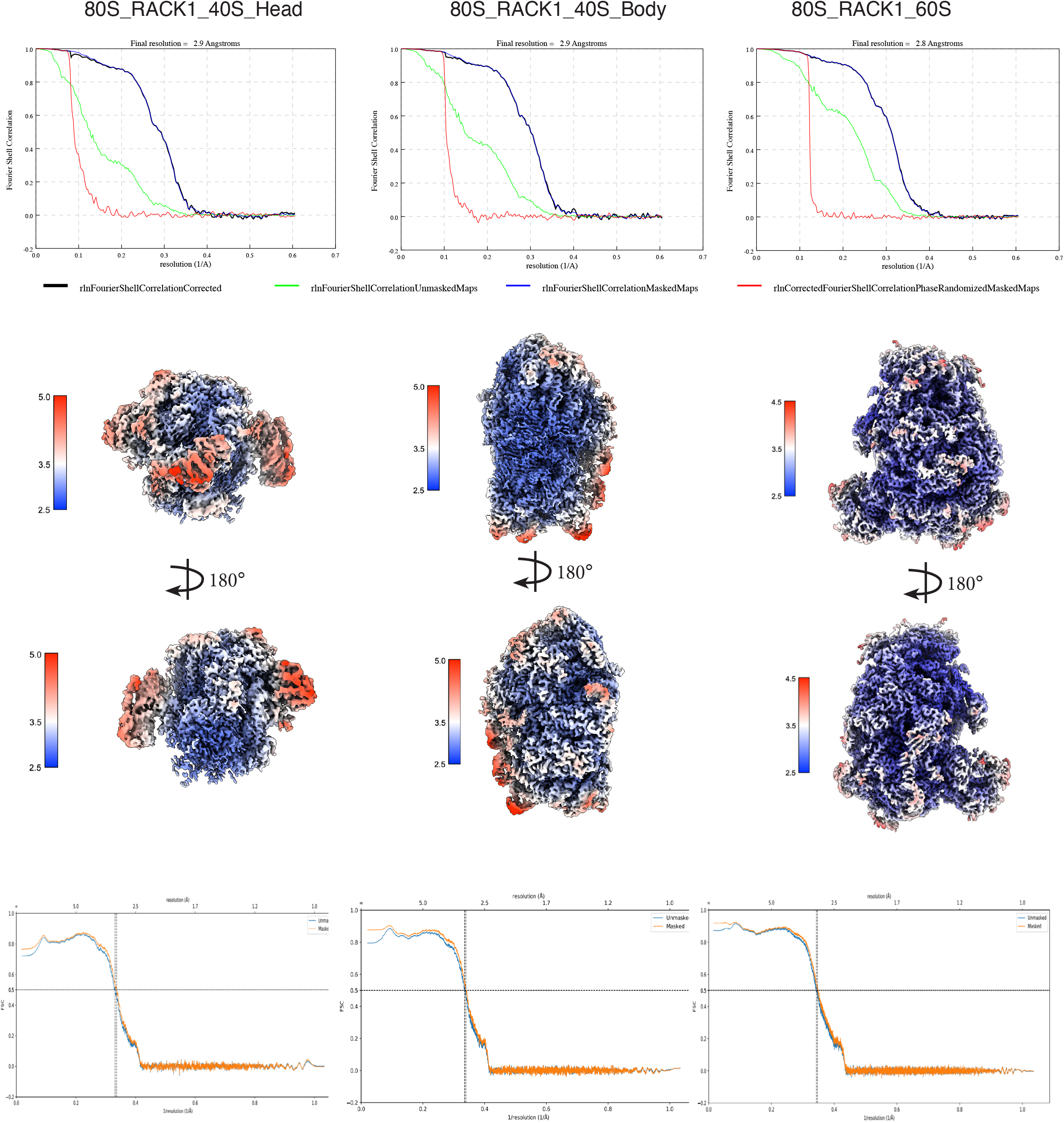
*T. gondii* 80S RACK1 local resolution information of postprocess maps and atomic models. **(A)**. Final resolution/FSC curve of density maps for 40S head, 40S body and 60S respectively; **(B)**. local resolution information of density maps for 40S head, 40S body and 60S respectively; **(C)**. Final resolution/FSC curve of atomic models for 40S head, 40S body and 60S respectively

**Supplemental Figure 5:**
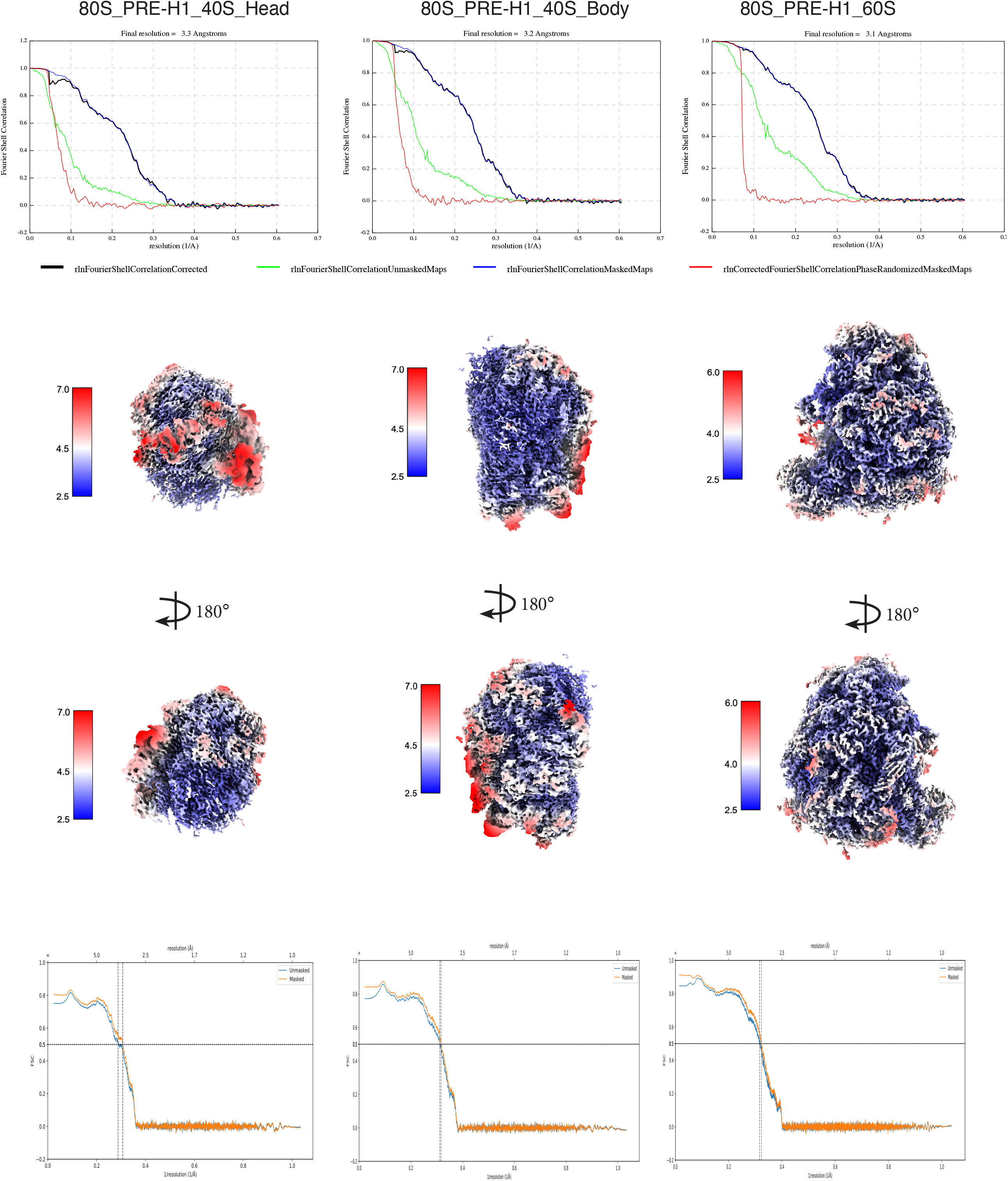
*T. gondii* 80S PRE-H1 local resolution information of postprocess maps and atomic models. **(A)**. Final resolution/FSC curve of density maps for 40S head, 40S body and 60S respectively; **(B)**. local resolution information of density maps for 40S head, 40S body and 60S respectively; **(C)**. Final resolution/FSC curve of atomic models for 40S head, 40S body and 60S respectively.

**Supplemental Figure 6:**
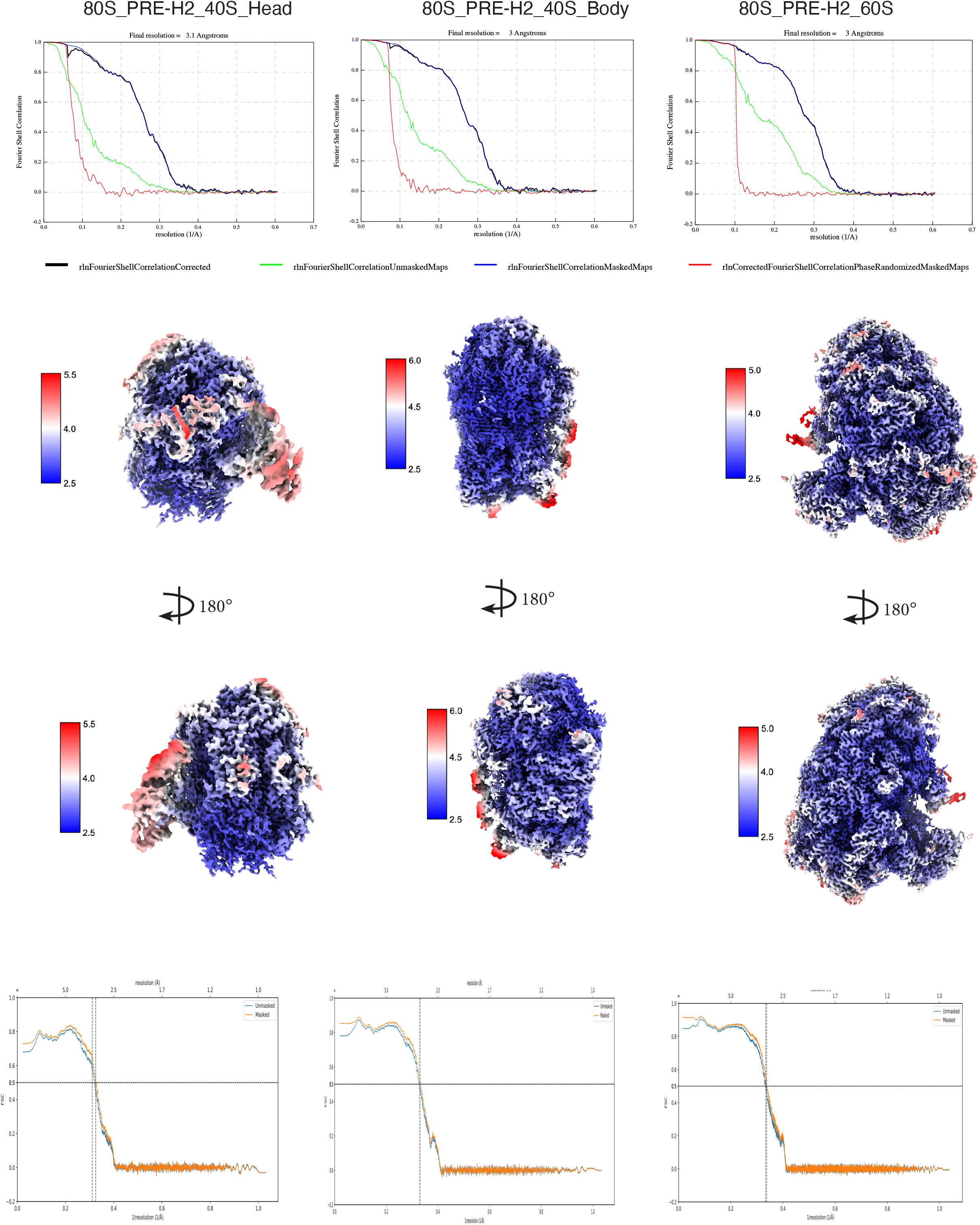
*T. gondii* 80S PRE-H2 local resolution information of postprocess maps and atomic models. **(A)**. Final resolution/FSC curve of density maps for 40S head, 40S body and 60S respectively; **(B)**. local resolution information of density maps for 40S head, 40S body and 60S respectively; **(C)**. Final resolution/FSC curve of atomic models for 40S head, 40S body and 60S respectively.

**Supplemental Figure 7:**
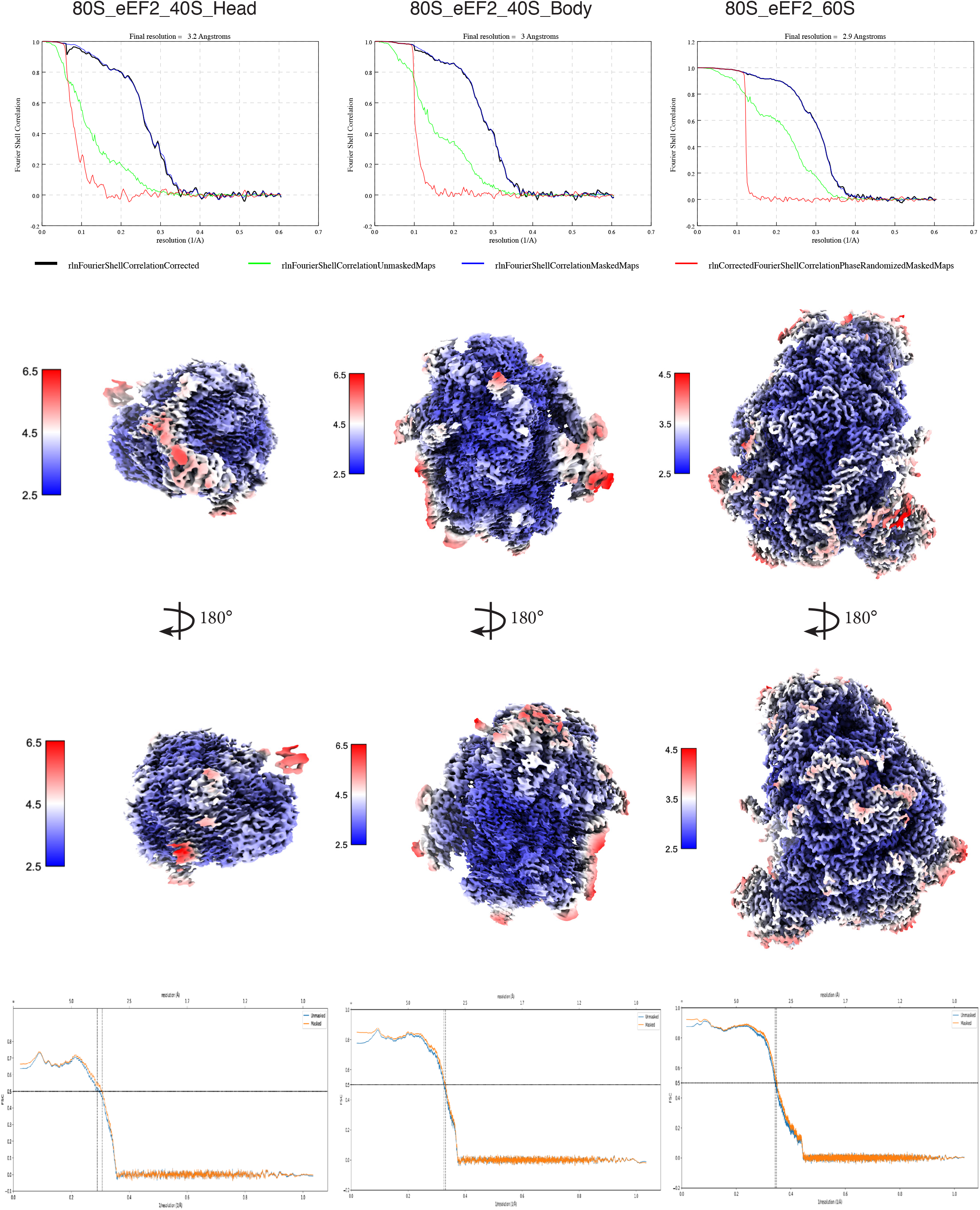
*T. gondii* 80S eEF2 local resolution information of postprocess maps and atomic models. **(A)**. Final resolution/FSC curve of density maps for 40S head, 40S body and 60S respectively; **(B)**. local resolution information of density maps for 40S head, 40S body and 60S respectively; **(C)**. Final resolution/FSC curve of atomic models for 40S head, 40S body and 60S respectively.

**Supplemental Figure 8:**
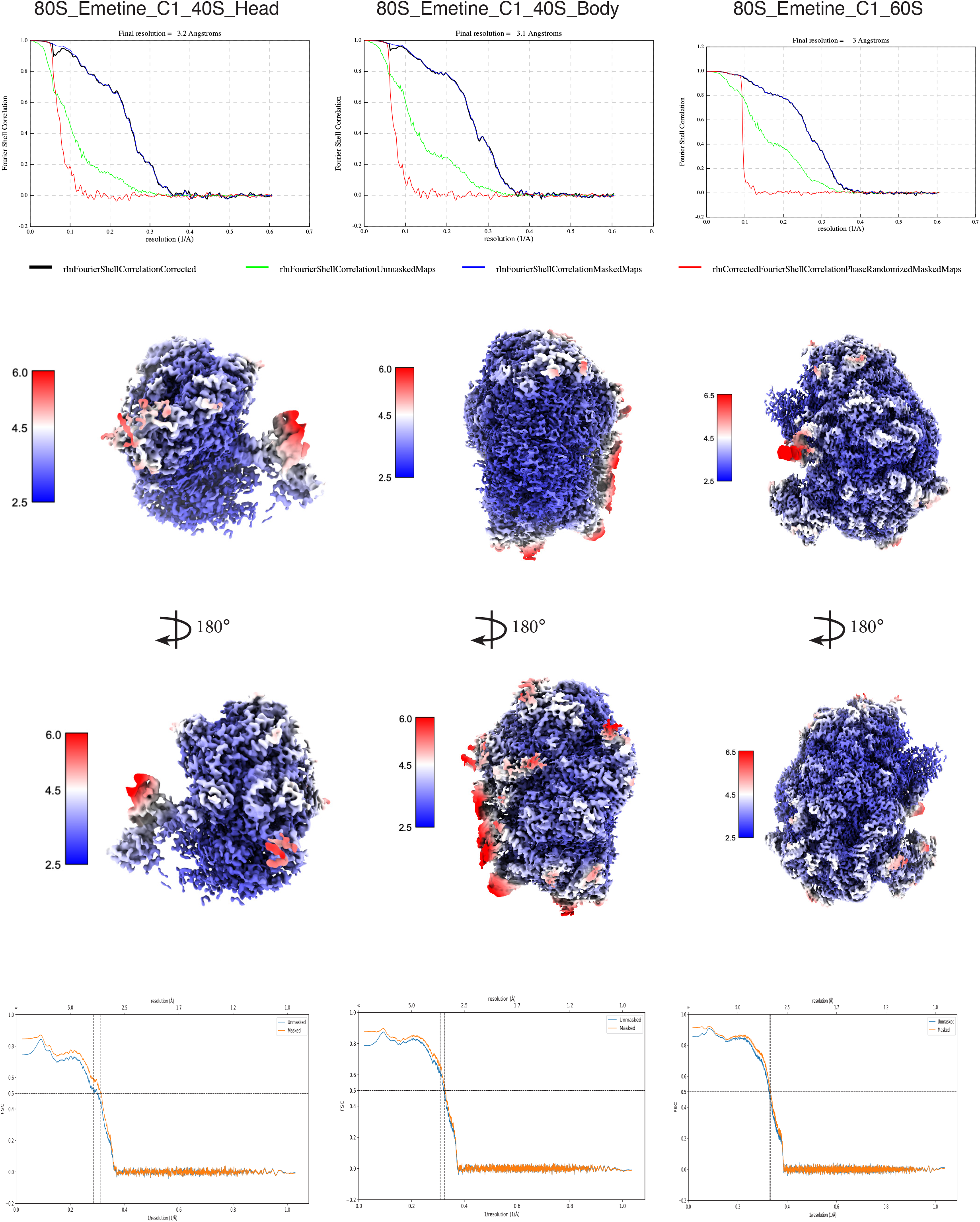
*T. gondii* 80S Emetine Class1 local resolution information of postprocess maps and atomic models. **(A)**. Final resolution/FSC curve of density maps for 40S head, 40S body and 60S respectively; **(B)**. local resolution information of density maps for 40S head, 40S body and 60S respectively; **(C)**. Final resolution/FSC curve of atomic models for 40S head, 40S body and 60S respectively

**Supplemental Figure 9:**
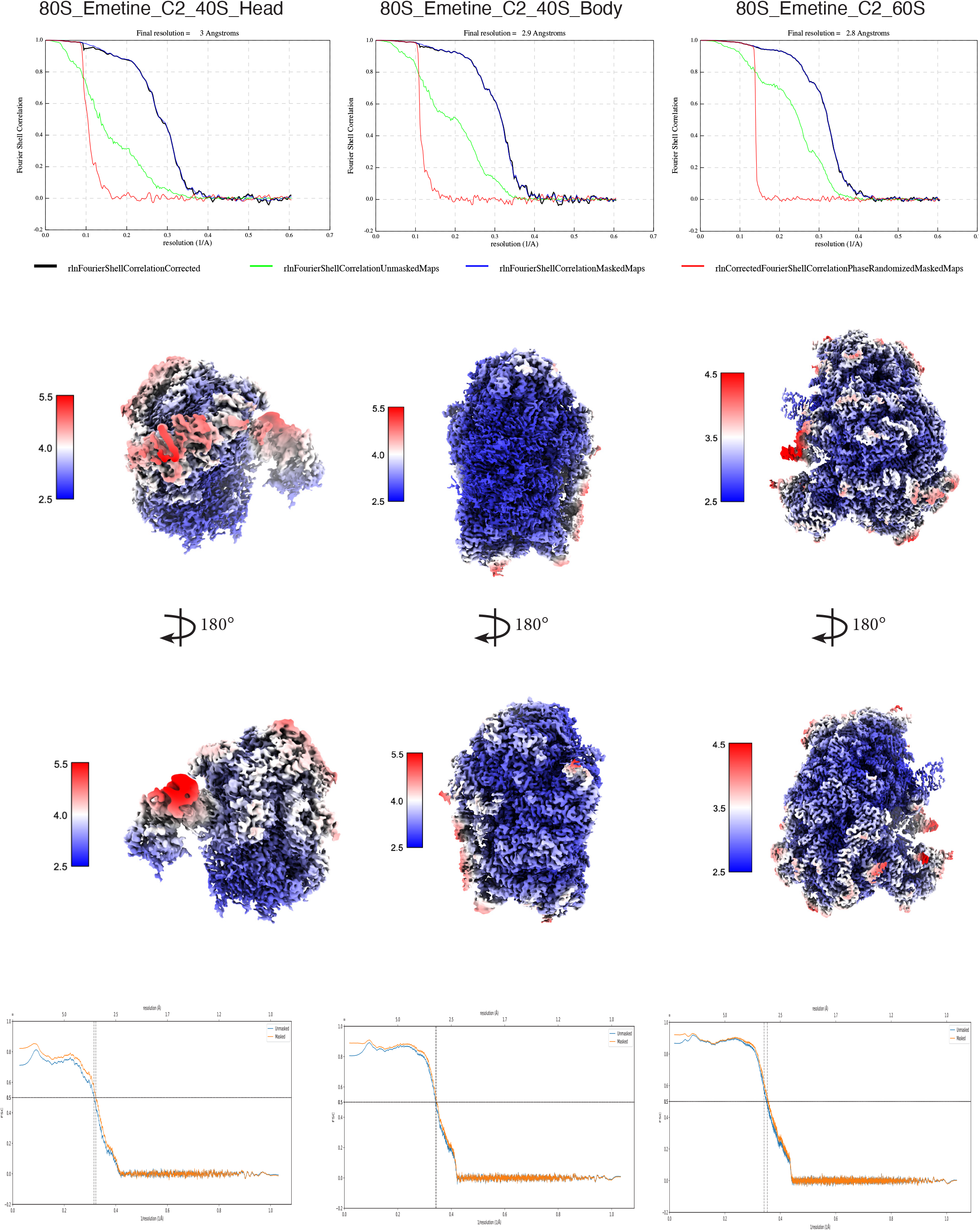
*T. gondii* 80S Emetine Class2 local resolution information of postprocess maps and atomic models. **(A)**. Final resolution/FSC curve of density maps for 40S head, 40S body and 60S respectively; **(B)**. local resolution information of density maps for 40S head, 40S body and 60S respectively; **(C)**. Final resolution/FSC curve of atomic models for 40S head, 40S body and 60S respectively

**Supplemental Figure 10:**
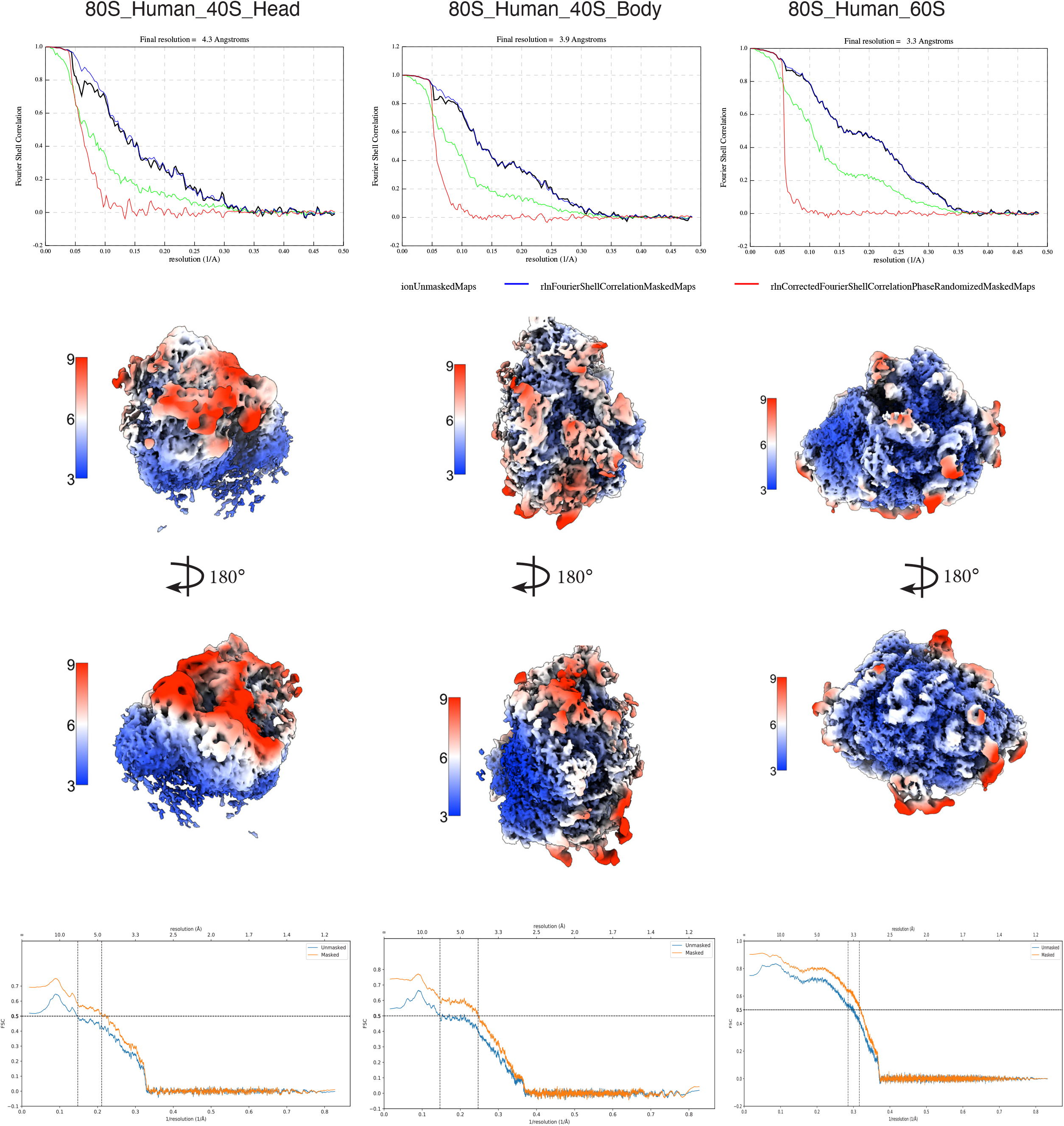
Human 80S local resolution information of postprocess maps and atomic Models. **(A)**. Final resolution/FSC curve of density maps for 40S head, 40S body and 60S respectively; **(B)**. local resolution information of density maps for 40S head, 40S body and 60S respectively; **(C)**. Final resolution/FSC curve of atomic models for 40S head, 40S body and 60S respectively

**Supplemental Figure 11:**
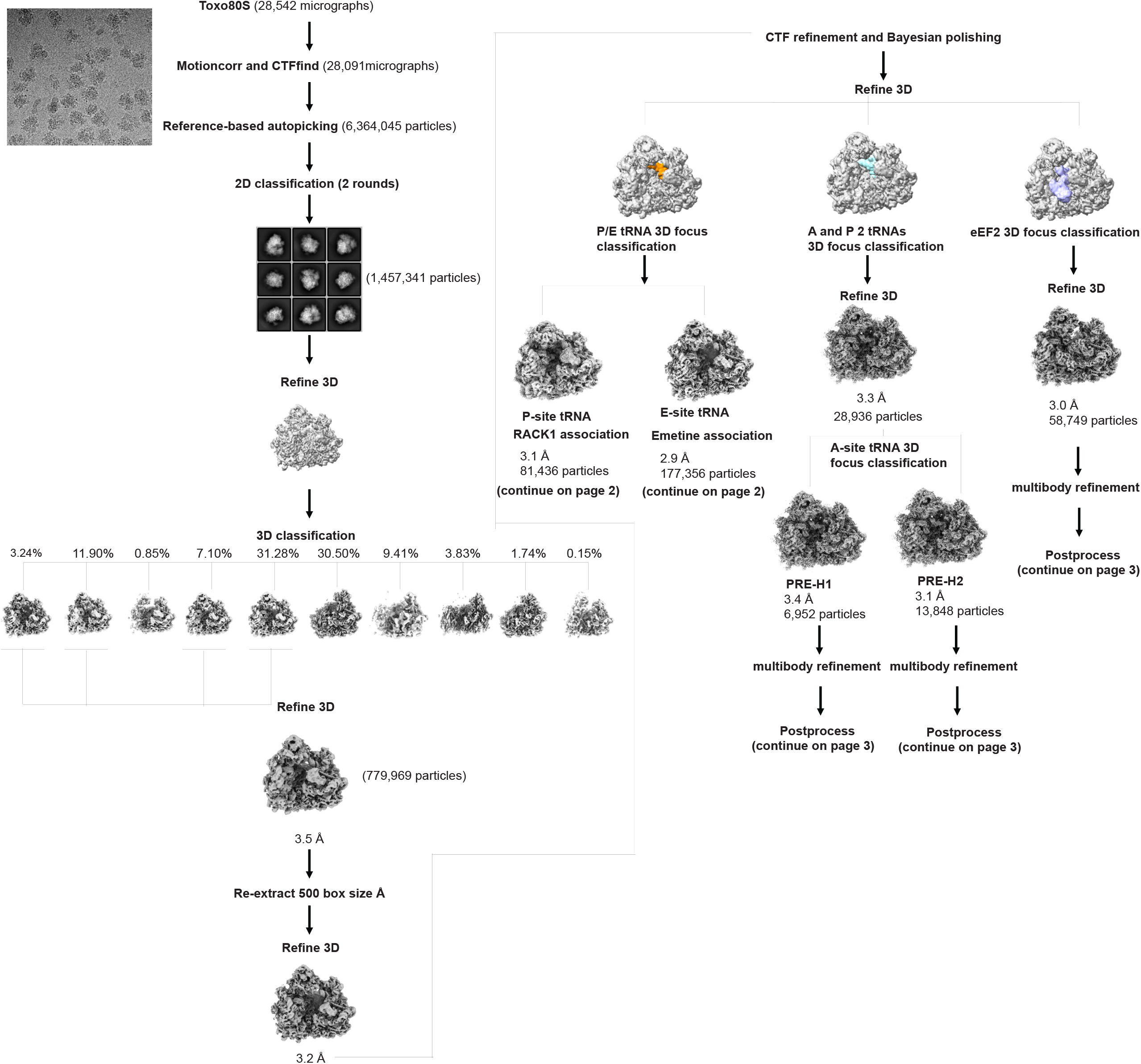
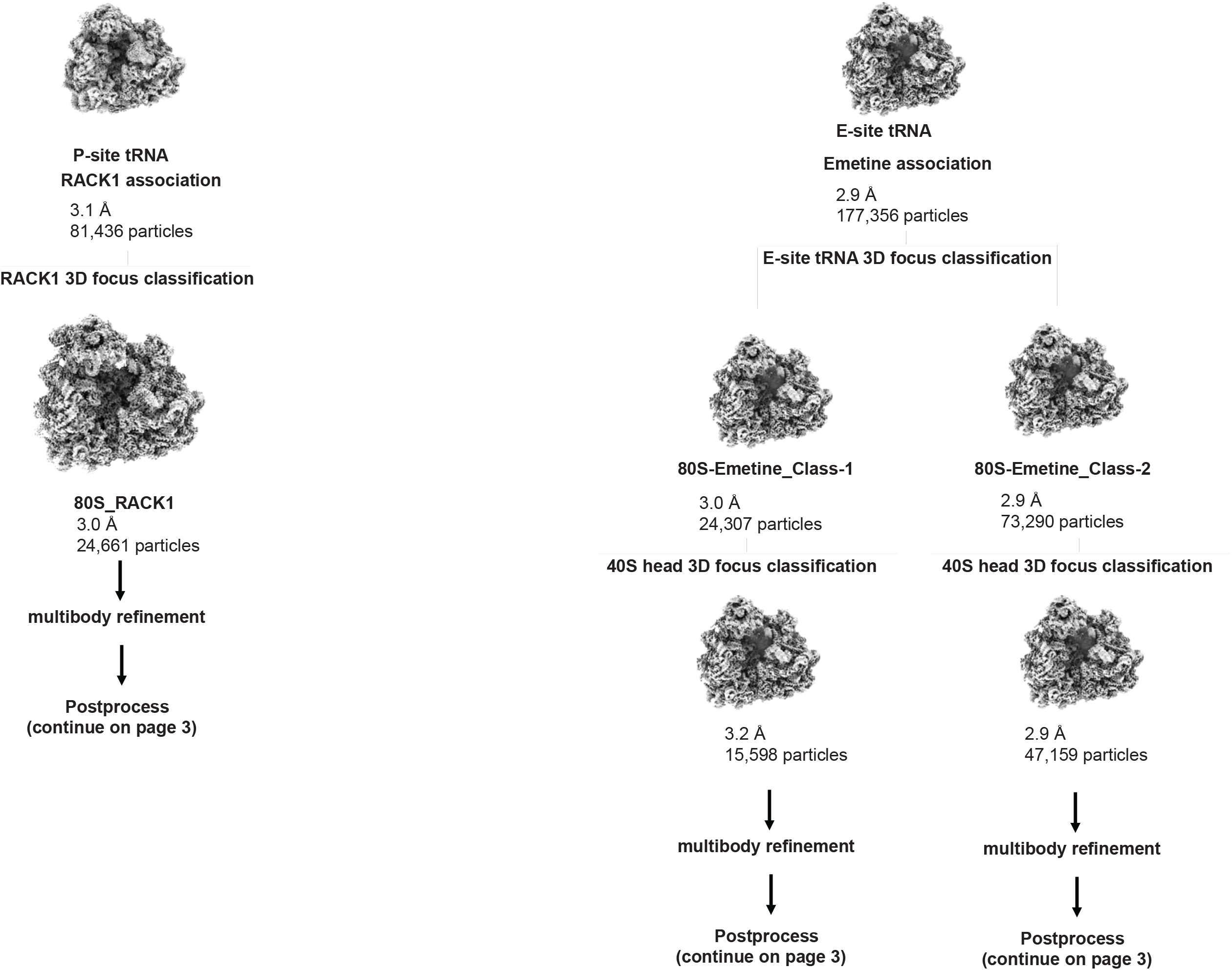
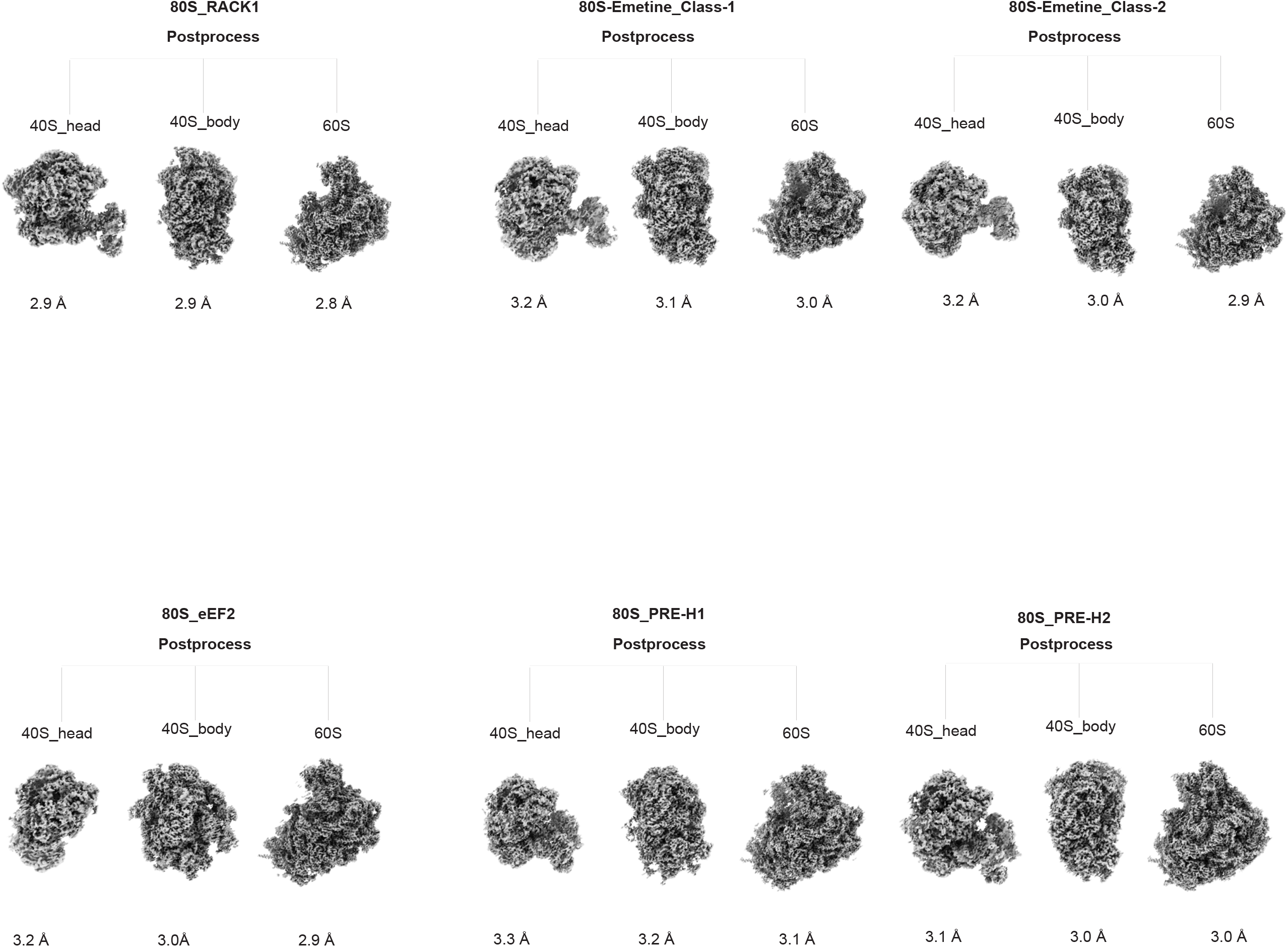
*T. gondii* 80S data process flow chart.

**Supplemental Figure 12:**
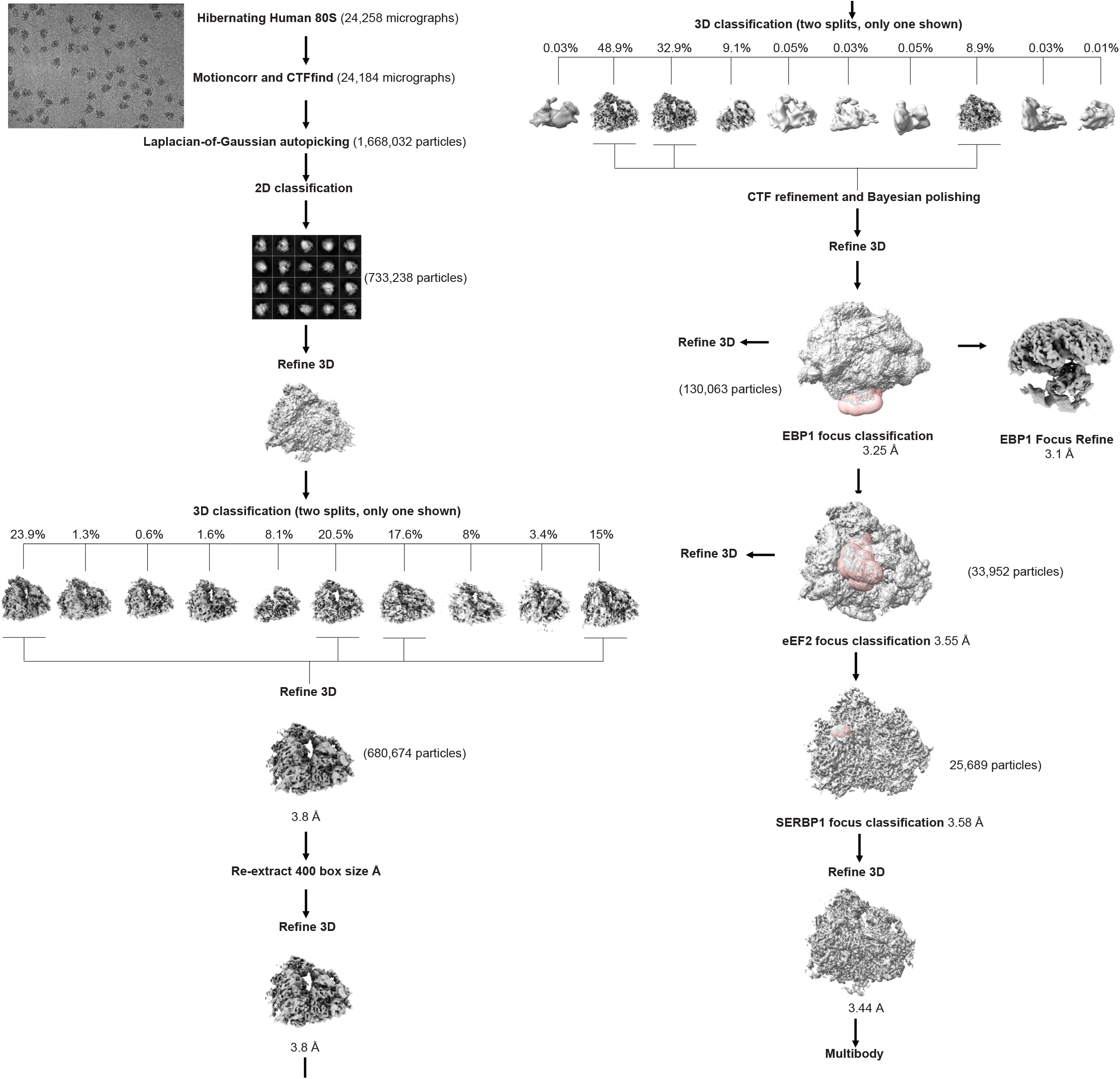
Human 80S data process flow chart.

**Supplementary Table 1.** Mass spectrometry analysis of purified *T. gondii* 80S ribosome complexes. LC–MS/MS analysis of proteins identified in purified *T. gondii* 80S ribosome complexes. The table includes identified proteins and peptides, sequence coverage, peptide-spectrum matches (PSMs), quantitative abundance values, and functional annotations generated from proteomic analysis.

**Supplementary Table 2.**
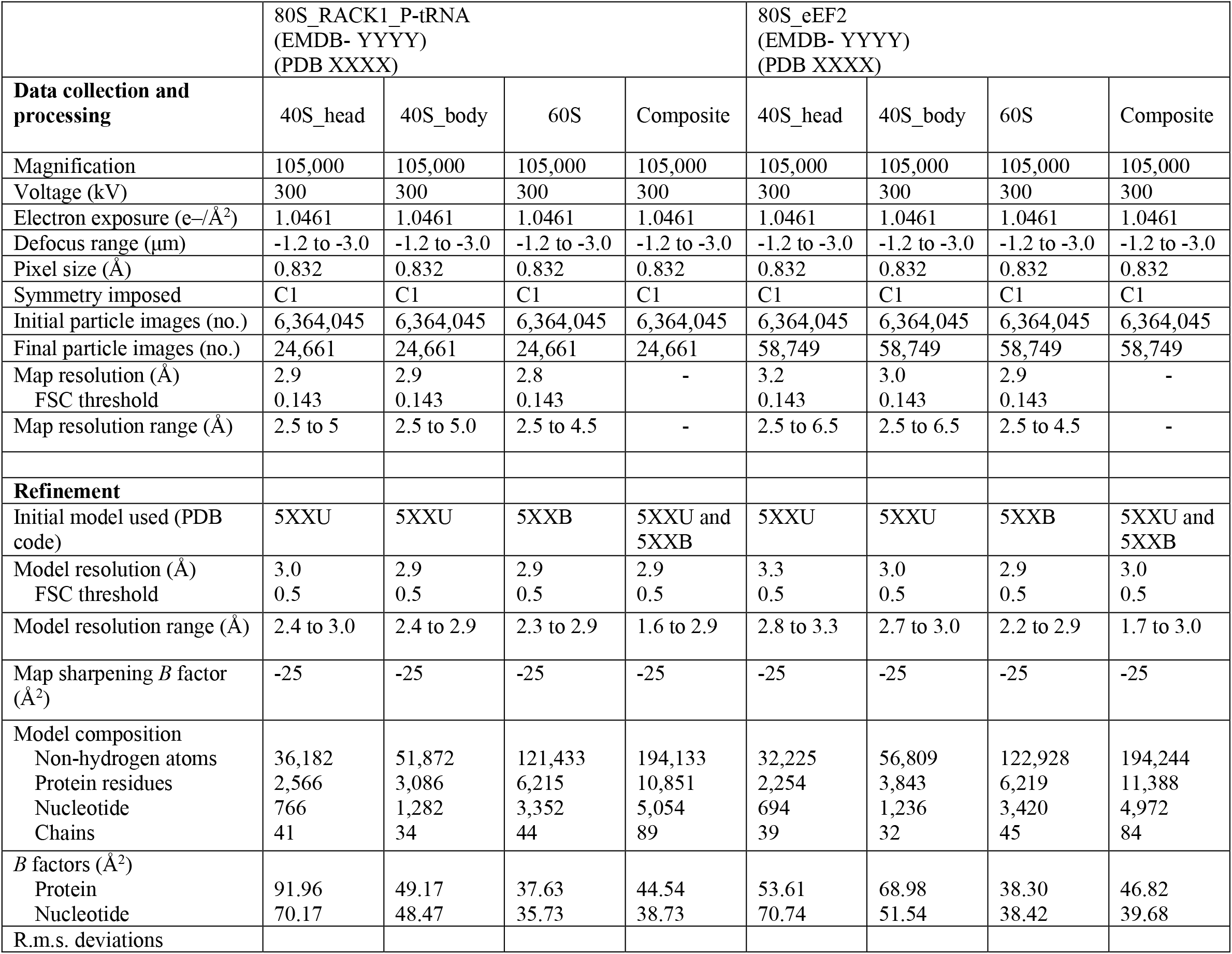
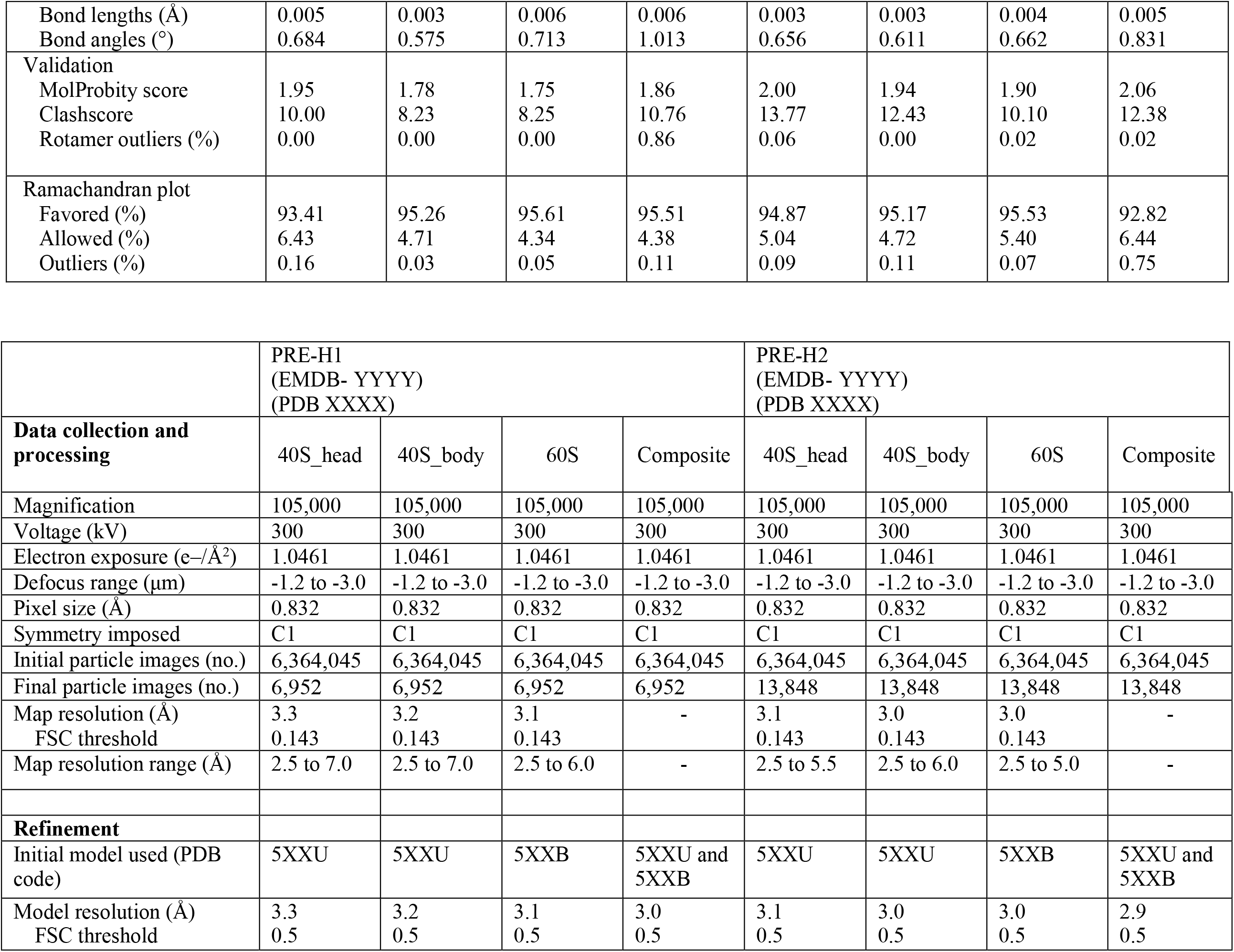
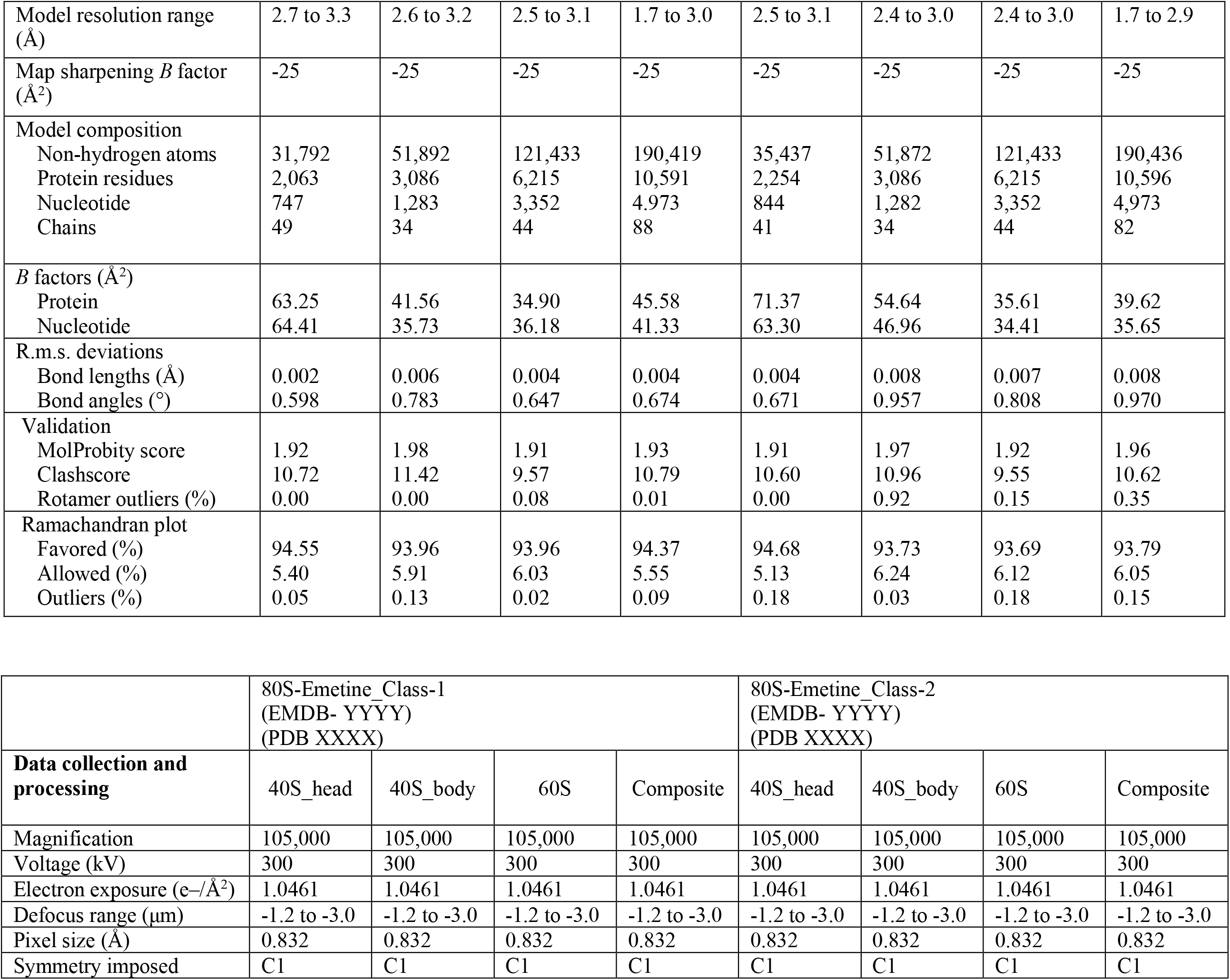
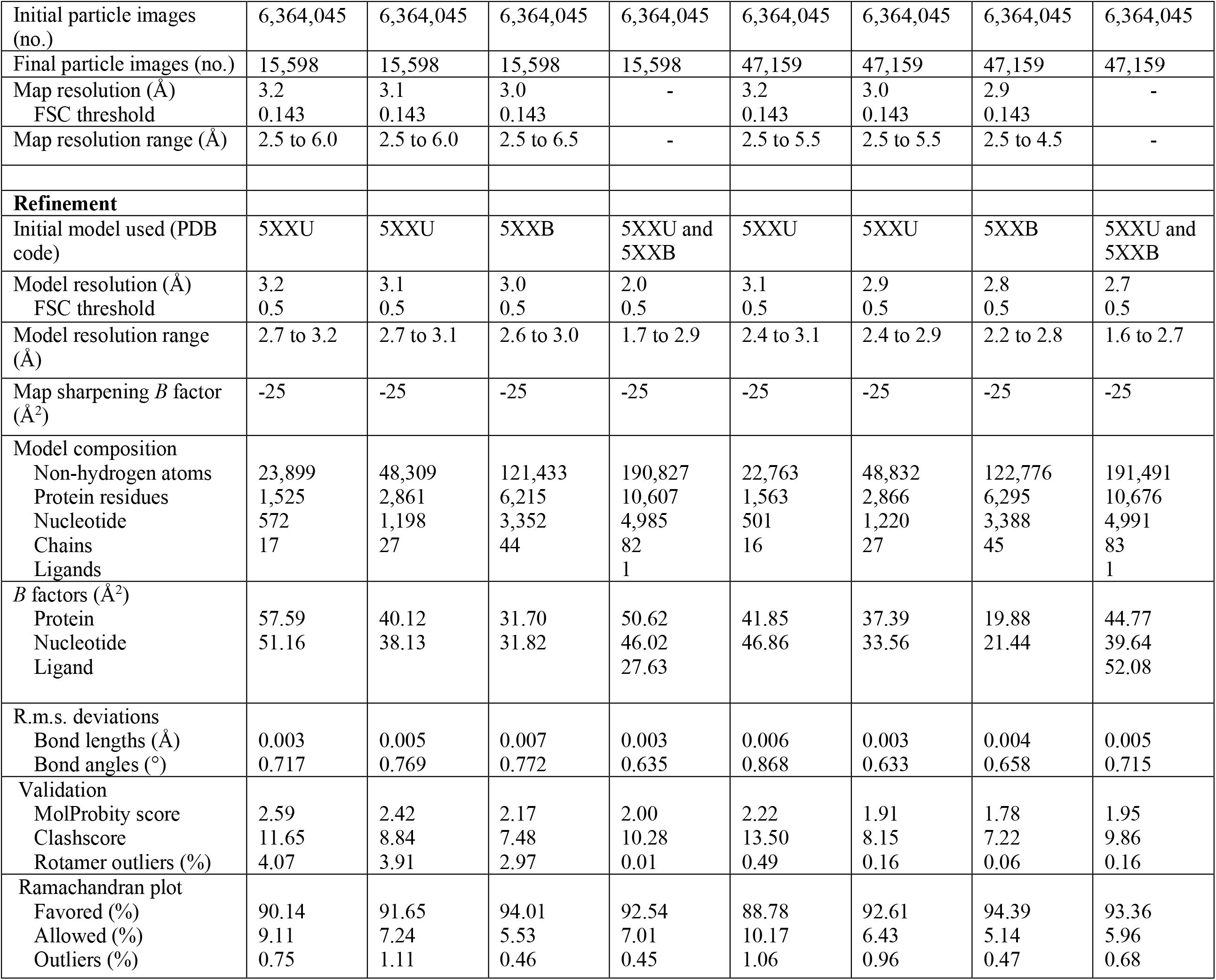
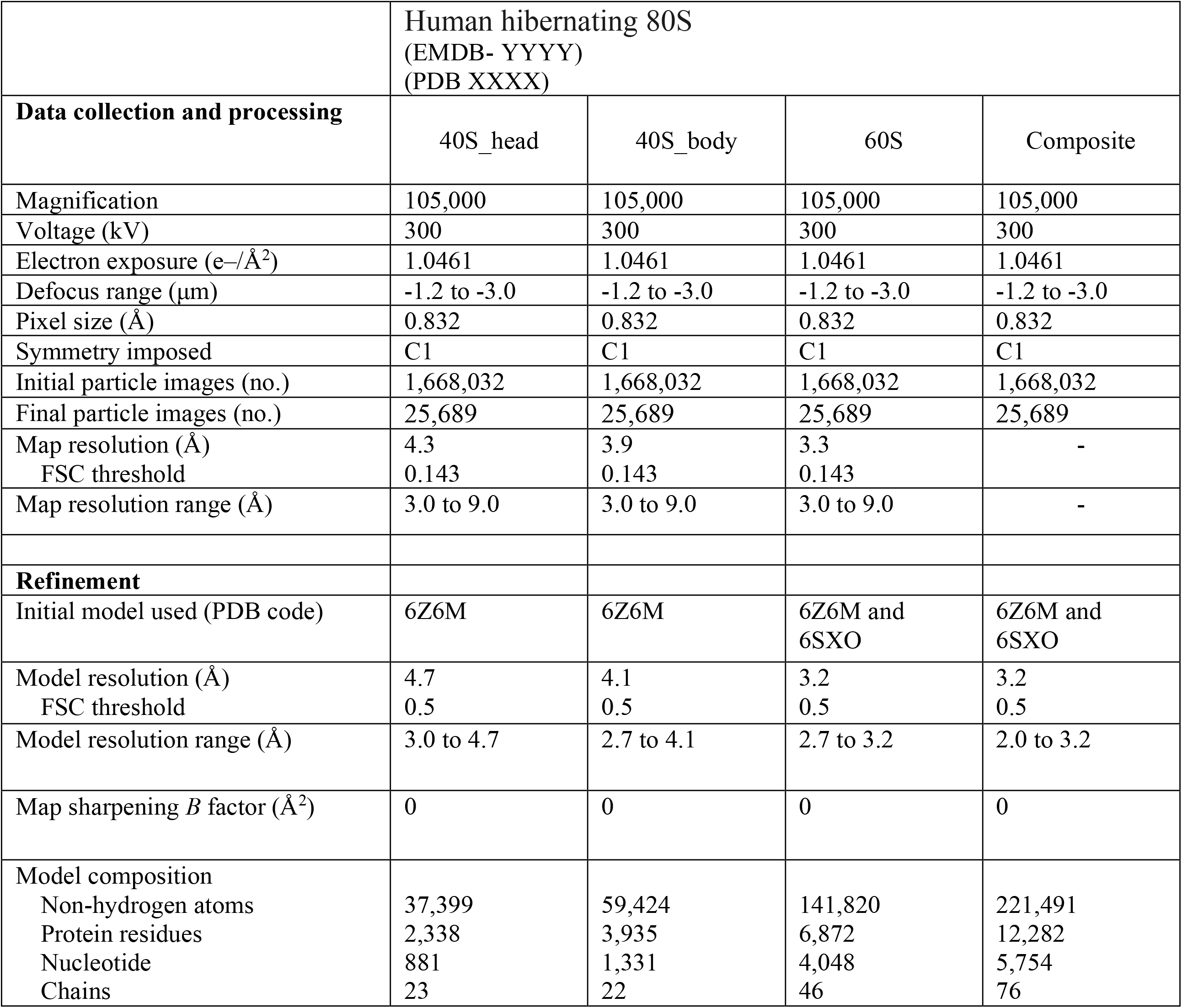
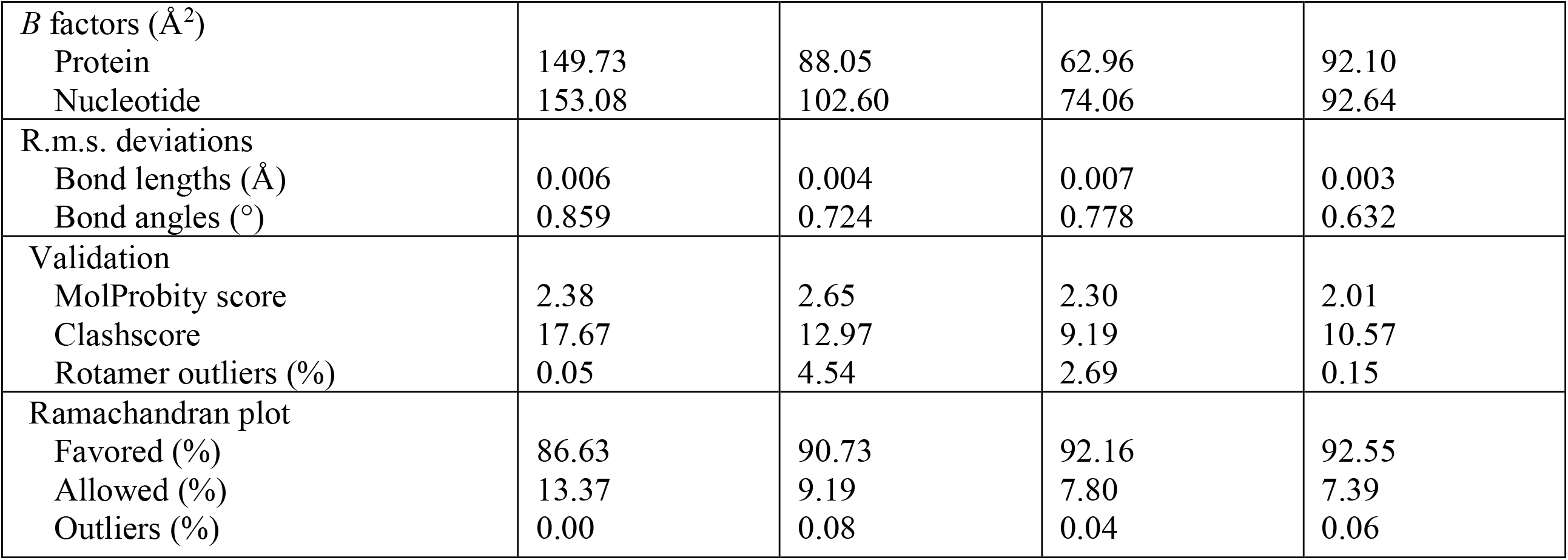
Cryo-EM data collection, refinement and validation statistics.

